# A Comparison of Brain Metabolic Connectivity Methods: Topology, Cognition, Age, Structure, and Genetics

**DOI:** 10.64898/2026.09.16.752264

**Authors:** Hamish A. Deery, Shenyue Zhao, Emma X. Liang, Rui Zheng, Sharna D. Jamadar

## Abstract

Brain metabolic connectivity is increasingly studied using positron emission tomography (PET), but multiple analytical approaches exist with unknown comparability. Here we compared metabolic connectivity approaches in 80 participants who underwent simultaneous dynamic PET with ^18^F-fluorodeoxyglucose and functional magnetic resonance imaging (fMRI) scans and cognitive testing. Six connectivity approaches were compared, including the traditional across-subject method (often labelled ‘metabolic covariance’) and five approaches that estimate individual-level connectivity: across-subject metabolic covariance with individual scores derived using leave one out (LOO); functional PET (fPET) within-subject correlations of regional timeseries; and three approaches that use distance-based measures to estimate individual-level connectivity based on within-region tracer distribution and kinetic compartment C1 and C2 time activity curves. fMRI connectivity was also calculated. Each method showed distinct topological, behavioural, structural and genetic signatures. fPET connectivity demonstrated the strongest cognitive associations and largest age-related declines, with unique predictive power for cognition independent of age. Tracer distribution showed strong age group discrimination although with mixed direction of age and cognitive effects. Metabolic covariance (LOO) and kinetic C2 showed no significant age group differences and their cognition associations reflected age-related variance. Gene expression associations implicated a shared core glycolytic-insulin machinery in all approaches, but the direction of the association with regional connectivity diverged across approaches and unique gene signatures further distinguished most methods. Collectively, these results indicate that the optimal connectivity approach depends on the research question. fPET connectivity with dynamic scans offers utility for cognition and ageing studies, tracer distribution for single-scan designs with careful interpretation of age and cognition relationships, kinetic approaches for novel biological insights into glucose kinetics and fMRI for multi-modal integration. Metabolic covariance (LOO) should be used with caution for brain-behaviour investigations. Researchers should weigh trade-offs between physiological specificity, analytical complexity, acquisition requirements and interpretability when selecting a metabolic connectivity approach.

## 1. Introduction

The study of brain networks has provided ground-breaking knowledge about the organisation and function of the human brain. Over the past two decades, the conceptualisation of the brain as a complex network of interconnected regions, the *connectome* (Sporns et al., 2005), has fundamentally reshaped our understanding of how cognitive processes emerge from distributed neural interactions. Functional and structural connectivity approaches, particularly those derived from functional magnetic resonance imaging (fMRI), electrophysiology (EEG) and diffusion-weighted imaging (DWI), have revealed that brain networks exhibit an efficient small-world topology (Bassett & Bullmore, 2006), contain highly connected hub regions (van den Heuvel & Sporns, 2011), and show reproducible large-scale organisational principles that are disrupted in normative ageing and neurological and psychiatric conditions (Deery et al., 2023b; Elam et al., 2021; Fornito et al., 2015; He et al., 2019; Preti et al., 2017; van den Heuvel & Hulshoff Pol, 2010). These insights have established connectomics as a cornerstone of modern neuroscience, providing a framework for understanding brain network organisation and linking brain architecture to behaviour, cognition, ageing and disease.

Although fMRI-based functional connectivity has been a dominant modality, the study of the connectome using molecular imaging actually predates fMRI, with the first study using ^18^F-fluorodeoxyglucose (FDG) positron emission tomography (PET) reported over 40 years ago (Horwitz et al., 1984). More recently, there has been a resurgence of interest in metabolic brain networks, driven by advances in FDG-PET imaging and a growing recognition that brain function is fundamentally constrained by glucose metabolism (Jamadar, 2026; Sala et al., 2023; Tuan et al., 2026; Yakushev et al., 2017). Compared with fMRI, which measures the haemodynamic response as an indirect proxy of neural activity, FDG-PET offers the advantage of directly indexing glucose metabolism, the primary energy substrate of the brain (Jamadar et al., 2025; Jamadar, Ward, et al., 2021). Furthermore, the emergence of functional PET (fPET) has enabled the measurement of temporal dynamics of glucose metabolism within individuals, opening new avenues for investigating metabolic connectivity and the time-variance of glucose signals with timescales approaching those of fMRI (Deery, Liang, Siddiqui, et al., 2024; Godbersen et al., 2023; Hahn et al., 2024; Jamadar et al., 2019; Jamadar, Ward, et al., 2021; Villien et al., 2014).

However, with this renewed interest in PET-derived connectivity has come a growth in diverse analytical approaches (reviewed in (Jamadar, 2026)), without a clear understanding of their relative strengths, weaknesses and the extent to which they capture similar or distinct aspects of brain organisation. These approaches differ fundamentally in their underlying data and computational methods (Jamadar, Ward, et al., 2021; Reed, Cocchi, et al., 2025) (Figure 1). The traditional approach calculates across-subject covariance of regional uptake values (Horwitz et al., 1984) and estimates no individual-level parameters (hereafter, ‘Metabolic Covariance - Across-Subjects’). Other methods derive individual-level connectivity measures. One derives individual-level parameters from the across-subject metabolic covariance matrix using leave-one-out (LOO) methods (hereafter referred to as ‘Metabolic Covariance (LOO)’) (Huang et al., 2020; Sun et al., 2022; Titov et al., 2017; Titov et al., 2014; Yao et al., 2018). Another derives connectivity from within-subject temporal correlations of fPET timeseries (Deery, Liang, Siddiqui, et al., 2024; Voigt et al., 2023) ( ‘fPET Connectivity’). More recent approaches capture voxel-wise tracer distribution patterns (Labarthe et al., 2026) via pairwise metabolic distances between region distributions (‘Tracer Distribution Connectivity’), or model the kinetic compartments of FDG uptake, namely, a delivery-like component to index the free intracellular tracer compartment (C1) and a trapping-like component to index the phosphorylated tracer component (C2), with networks derived from kinetic similarity of the regional time activity curves (TACs) (Facca et al., 2025; Facca et al., 2026; Volpi et al., 2023) (‘Kinetic C1’ and ‘Kinetic C2 Connectivity’, respectively). Each of these methods potentially reveals different facets of brain metabolism, yet they have not been systematically compared within the same cohort.

**Figure 1.**
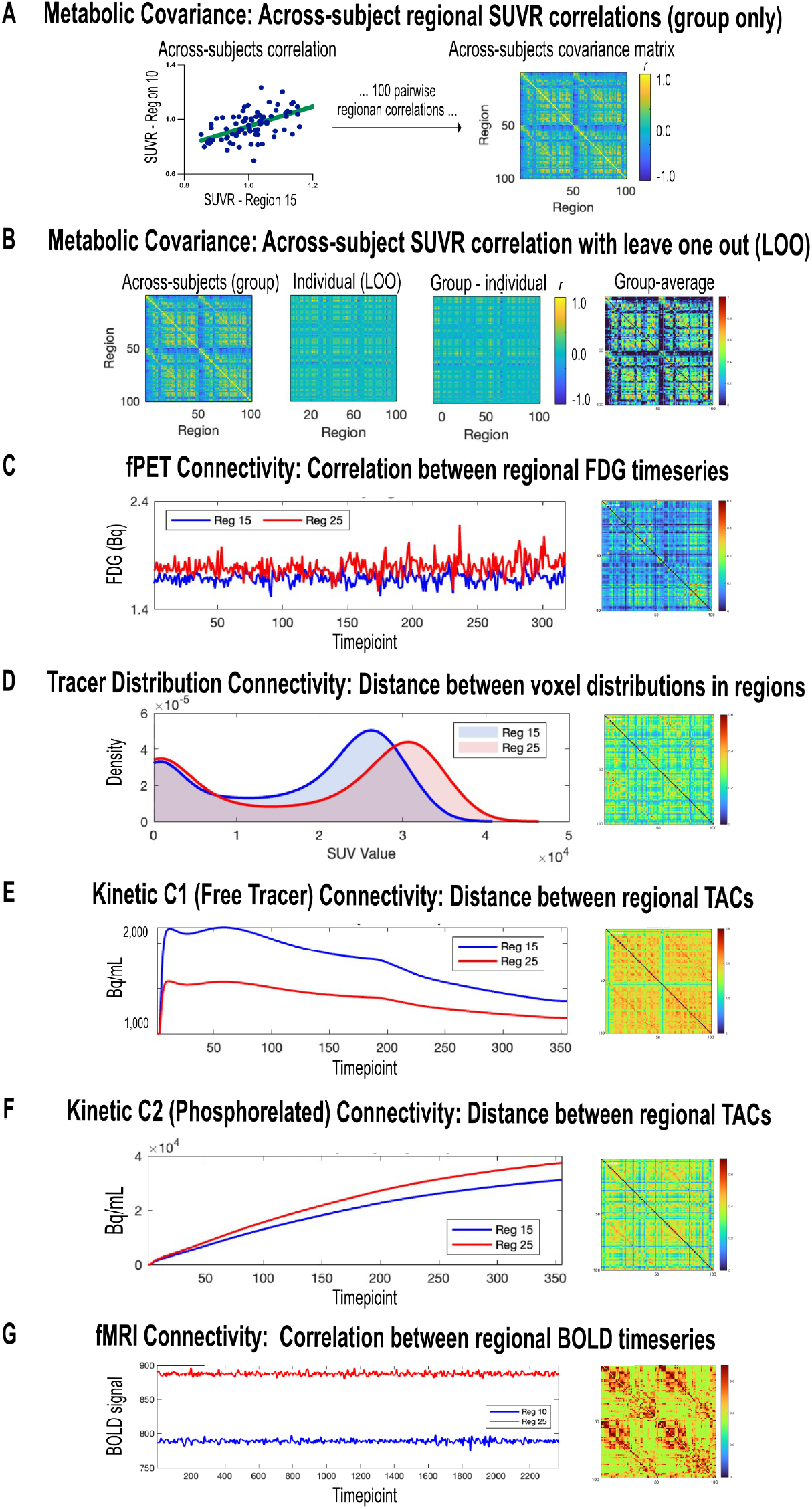
Overview of connectivity methods. Illustration of representative regional data and connectivity matrices from each approach. Connectivity was estimated as: (A) Metabolic Covariance, derived from across-subject covariance of regional SUVR values at the group level only; (B) Metabolic Covariance, derived from across-subject covariance of regional SUVR values at the group level, with subject-specific contributions estimated via leave-one-out (LOO); (C) fPET connectivity, derived from within-subject temporal correlations of FDG time-activity curves; (D) Tracer Distribution connectivity, calculated from voxel-wise distribution similarity of static FDG uptake within regions; (E) Kinetic C1 Connectivity, derived from Euclidean similarity of free intracellular tracer concentration time-activity curves; (F) Kinetic C2 Connectivity, derived from Euclidean similarity of phosphorylated tracer (¹⁸F-FDG-6P) concentration time-activity curves; and (G) fMRI Connectivity, derived from within-subject temporal correlations of BOLD timeseries. All connectivity matrices were parcellated into 100 cortical regions using the Schaefer100 atlas.

Each metabolic connectivity method also requires different considerations of the underlying biological processes that they measure. fPET Connectivity captures the temporal co-fluctuations of glucose uptake across brain regions (Jamadar, Ward, et al., 2021). This approach provides a view of dynamic metabolic coupling in the temporal domain, analogous to EEG and fMRI functional connectivity but grounded in glucose metabolism. Recent evidence suggests that fPET Connectivity provides unique and complementary information about brain networks relative to fMRI and that the temporal dynamics of the underlying fPET signal carries biologically meaningful information about network efficiency, state switching, cognition and ageing (Deery et al., 2025a; Deery, Liang, et al., 2026; Deery et al., 2025b; Deery, Liang, Siddiqui, et al., 2024). Metabolic Covariance (Across-Subject), in contrast, is estimated from across-subject correlations of regional standardised uptake value ratios (SUVR), and therefore reflects commonalities in regional uptake across subjects but may not reflect underlying subject-level responses due to statistical ergodicity (Jamadar, Ward, et al., 2021; Medaglia et al., 2011). However, Metabolic Covariance (LOO) can be used to derive subject-specific results (Yao et al., 2018). Although the biological and functional relevance of across subject covariance methods has been questioned (Jamadar, Ward, et al., 2021; Reed, Cocchi, et al., 2025; Sala et al., 2022; Volpi et al., 2023), both approaches (Across-Subject and LOO) have been used to study disease states, such as comparing people with neurological conditions to controls (Horwitz et al., 1984; Huang et al., 2020; Khokhar et al., 2023; Morbelli et al., 2012; Yao et al., 2018).

Tracer Distribution Connectivity is a recently developed method that quantifies similarity between regions based on their full voxel-wise distribution of FDG uptake ((Labarthe et al., 2026) also see (Li et al., 2023; Tuan et al., 2026; Wang et al., 2020) for related methods). Initial evidence indicates that the method has the ability to identify disease progression at the individual level when used in whole body (brain-body) PET studies (Labarthe et al., 2026). Finally, kinetic connectivity approaches derive TAC similarity from compartments C1 and C2 (Volpi et al., 2023). Hence, these kinetic measures directly quantify the underlying biochemistry of glucose metabolism, with recent reports that brain networks derived from these measures are altered in glioma (Vallini, Baron, et al., 2025; Vallini, Silvestri, et al., 2025).

One of the challenges of methodological comparison in brain connectivity is that there is no ground truth. Patterns of brain organisation are understood at a conceptual level (e.g., small-worldness, hubness), however, the true underlying patterns of connectivity are unknown. While fMRI connectivity benefits from a very large and diverse body of research and therefore serves as a useful comparison, it too cannot be used as ground truth, as it indexes related but fundamentally different physiology (Deery et al., 2025b; Jamadar, Ward, et al., 2021), with debatable reliability (Logothetis, 2008; Logothetis & Wandell, 2004). Therefore, a finding that each method differs in its topology and connectivity parameters can be difficult to evaluate. One useful model to compare methods is to test their ability to determine phenotypic variability, for example, differences by participant age, cognitive performance or other biological properties, such as anatomy and molecular spatial topology. Such an approach can provide information about the measurement validity of each method and yield insight into the applications or research questions that each method is best suited to address.

Given the diversity of metabolic connectivity approaches, limited previous comparisons, and the absence of ground truth, the present study had four aims. First, to characterise the network topology of six metabolic connectivity methods within the same cohort: Metabolic Covariance (Across-Subject), Metabolic Covariance (LOO), fPET Connectivity, Tracer Distribution Connectivity and Kinetic C1 and C2 Connectivity. We also examined fMRI Connectivity from simultaneously acquired data for an additional benchmark comparison. Note that because Metabolic Covariance (Across-Subject) provides no individual participant-level parameters, our subsequent aims were tested with the five PET connectivity approaches that provide individual parameters and fMRI. Our second aim was to examine how network topology varies across approaches and relates to age and cognitive performance. Third, we aimed to evaluate the alignment of each approach with structural connectivity. Fourth, we aimed to investigate the genetic basis of each approach by relating network measures to the expression of glucose metabolism genes. By systematically comparing these methods we aim to provide a comprehensive framework for characterising the networks and advancing our understanding of their relationship with brain structure and function. Understanding the relative strengths and limitations of each approach can also guide method selection for future studies.

## 2. Methods

### 2.1 Participants

The Monash *MetConn* simultaneous PET/MR dataset was used for the study. The dataset has been described in previous work (Deery, Liang, Di Paolo, et al., 2024; Deery et al., 2025b; Deery, Liang, Siddiqui, et al., 2024). Whole sample (N = 80) and younger (N = 36) and older (N = 44) age group differences in demographic characteristics and cognition are reported in Supplement 1. Briefly, the mean age of the whole sample was 54.5 years (SD = 24.5). The proportion of women was 50%. The average years of education was 17. Average BMI was 24.8 kg/m^2^, resting heart rate was 78.1 BPM and systolic and diastolic blood pressure were 136.2 and 78.1 mmHg, respectively. The mean fasting blood glucose was 5.0 mmol/L.

### 2.2 Data Acquisition: Cognitive Testing and Neuroimaging

Full details of the cognitive tests and neuroimaging scan parameters are provided in Supplement 2 and 3, respectively. Briefly, participants completed the Hopkins Verbal Learning Test (HVLT) (Shapiro et al., 1999), and computer-based task-switching (Friedman et al., 2008), stop-signal (Verbruggen et al., 2008), digit symbol substitution (Thorndike, 1919) and digit span (Blackburn & Benton, 1957) tests. Participants underwent a 90-minute simultaneous MR-PET scan in a Siemens Biograph 3-Tesla molecular MR scanner while watching a video of a drone flying over the Hawaii islands. The scan sequences included anatomical T1- and T2-weighted MRI scans, list-mode PET, and T2* EPI BOLD-fMRI and diffusion-weighted (DWI) sequences. At the start of the scan, half of the 260 MBq FDG tracer was administered as a bolus with the remainder infused over 50 minutes at 36 mL/hour (Jamadar et al., 2019; Jamadar et al., 2020).

### 2.3 Neuroimaging Pre-processing

Full pre-processing details are provided in Supplement 3. Briefly, PET data were binned into 344 frames (16s each), attenuation-corrected (Burgos et al., 2014) and reconstructed using OSEM with point spread function correction. Reconstructed volumes were converted to NIFTI format, assembled into 4D files, motion and partial volume corrected, and surface-based smoothed at 8mm FWHM (Greve et al., 2016; Greve et al., 2014). T2* images were unwarped, motion-corrected, detrended, and spatially smoothed at 8mm FWHM. The FDG and fMRI images were normalised to MNI space and their timeseries denoised using aCompCor (Whitfield-Gabrieli & Nieto-Castanon, 2012); for fPET this yielded residual metabolic connectivity time series capturing glucose signal dynamics while minimising the FDG uptake trend (Deery, Liang, et al., 2026; Deery, Liang, Siddiqui, et al., 2024; Reed, Ponce de Leon, et al., 2025). fMRI data were additionally bandpass filtered (0.01–0.1 Hz). Structural connectivity was derived from DWI using single-shell multi-tissue constrained spherical deconvolution and anatomically constrained probabilistic tractography, generating 22 million streamlines per participant with SIFT2 weighting (Dhollander, 2016; Smith et al., 2012, 2015; Tournier et al., 2019; Yeh, 2018).

### 2.4 Connectivity Measures

#### 2.4.1 Metabolic Covariance (Across-Subject and Leave One Out)

To derive Metabolic Covariance (Across-Subjects), dynamic PET frames from 30-60 minutes (Goutal et al., 2020) were averaged to create a single, static image and standard uptake values (SUVR) were extracted for each of the 100 Schaefer regions (Schaefer et al., 2018) by normalising to the whole-brain grey matter mean uptake. Metabolic Covariance (Across-Subject) was computed as the Pearson’s correlation between SUVR vectors across all subjects (R-all), yielding a 100 × 100 correlation matrix (Figure 1A). For Metabolic Covariance (LOO), this procedure was repeated with each subject sequentially removed: for subject *s*, the correlation matrix R-s was computed from the remaining n-1 subjects, and the LOO contribution for subject *s* was calculated as (n × R-all - (n - 1) × R-s) (Figure 1B).

#### 2.4.2 fPET Connectivity

For each participant, fPET Connectivity was computed as the Pearson’s correlation between all pairs of regional timeseries (Figure 1C). The first 10 minutes of the scan were excluded to minimise any post-denoising residual impact on connectivity estimates of the initial FDG uptake (Deery et al., 2025b; Deery, Liang, Siddiqui, et al., 2024). The correlation matrix was Fisher z-transformed and saved for each subject.

#### 2.4.3 Tracer Distribution Connectivity

To measure Tracer Distribution Connectivity, we adapted a recently developed framework for individualised PET analysis from single scan protocols (Labarthe et al., 2026) (Figure 1D and Supplement 4 for details). The average PET images was calculated from the 10-minute point onwards, from which voxel-wise uptake distributions were extracted for each of the 100 cortical regions. To reduce dimensionality while maintaining distributional shape, a spatially aware compression algorithm was used. Connectivity between regions was computed using energy distance between voxel intensity distributions, corrected for non-linear dependencies via mutual information. The final connectivity strength was computed using an exponential decay function, emphasising strong distributional similarities while down-weighting distant or independent relationships.

#### 2.4.4 Kinetic Connectivity (C1 and C2)

To obtain physiologically meaningful kinetic connectivity measures, we extracted the TACs from the dynamic FDG-PET data, closely following the approach developed by Castellaro and colleagues (Castellaro et al., 2017) and used by Volpi et al. for connectivity analysis (Volpi et al., 2023) (Figure 1E–F; Supplement 4 for details). Briefly, we estimated voxel-wise parameters of Sokoloff’s two-tissue, three-rate-constant model using variational Bayesian inference, yielding parametric maps of tracer inflow, efflux, phosphorylation, and blood volume. From these, region-level time-activity curves (TACs) were constructed for two components, a delivery-like component and a trapping-like component. These two components sum to the total fitted tissue curve and provide complementary time courses separating delivery-related from trapping-related information, echoing the C₁ and C₂ distinction of Sokoloff’s model. For each subject, connectivity between brain regions was computed as the Euclidean distance between regional TACs of each component, normalised to a similarity scale (0-1) and Fisher z-transformed.

#### 2.4.5 fMRI Connectivity

fMRI Connectivity was computed as the Pearson’s correlation between all pairs of regional timeseries (Figure 1G), with the correlation matrix Fisher z-transformed for each subject.

#### 2.4.6 Structural Connectivity

The structural connectivity matrix was constructed with each element representing the strength of the connection between a pair of regions, calculated as the sum of SIFT2-weighted streamlines connecting them.

### 2.5 Gene Expression Data

Gene expression data were obtained from the Allen Human Brain Atlas (Shen et al., 2012) using the Abagen toolbox (Markello et al., 2021). Expression values were extracted for the 100 regions, as follows: (1) probe aggregation by selecting the probe with highest variance per gene, (2) mapping samples to regions using MNI centroids with affine transformation, (3) donor-independent processing with z-score normalisation, and (4) averaging across donors. Glucose metabolism genes were identified from Gene Ontology (GO:0006006), MSigDB Hallmark Glycolysis, KEGG Glycolysis, and glucose transporter gene lists, yielding 94 genes for analysis.

### 2.6 Statistical Analyses

#### 2.6.1 Sparsity Threshold Selection and Graph Metrics

To determine the optimal sparsity threshold for network construction, we employed the cost-efficiency framework introduced by Bassett and colleagues (Bassett et al., 2009). This approach balances the trade-off between network cost and global efficiency (Bassett & Bullmore, 2006; Bullmore & Sporns, 2012). This cost-efficiency analyses converged on a sparsity level of 15%-25% (see Supplement 5), with 15% applied uniformly across subsequent analyses. Sensitivity analyses at 25% sparsity are reported in Supplement 9.

For each connectivity approach, network topology was characterised using four graph metrics (Bullmore & Sporns, 2009; Sporns & Betzel, 2016): (1) Small-worldness (σ), measured as the ratio of normalised clustering to path length (> 1.0 indicate small-world organisation); and (2) rich-club coefficient (Φ), the extent to which high-degree nodes (hubs) are more densely interconnected than expected by chance. We also calculated two additional metrics to assist in understanding differences in small-worldness across approaches, as the additional metrics represent components of the small-world composite: (3) clustering coefficient (C), measures as the average fraction of a node’s neighbours that are also connected to each other, reflecting local network segregation; and (4) Characteristic path length (L) calculated as the average shortest path length between all pairs of nodes, reflecting global network integration. The formula for each graph metric is provided in Supplement 6.

#### 2.6.2 Network Topology Similarity by Connectivity Approach

To quantify the similarity between connectivity approaches, we computed DICE coefficients on group-average connectomes. To compare graph metrics across approaches, repeated-measures ANOVAs were performed with connectivity approach as the within-subjects factor and graph metric as the dependent variable. Post-hoc pairwise comparisons were Bonferroni-corrected for multiple comparisons across the 15 pairs of approaches.

#### 2.6.3 Age Group Differences by Connectivity Approach

Independent samples *t*-tests were used to compare younger (N=36, mean age 28.4 years) and older (N=44, mean age 75.8 years) adults on the graph metrics for each connectivity approach. To control for multiple comparisons, false discovery rate correction was applied at p-FDR < 0.05 for each approach. We also performed age group classification using quadratic discriminant analysis. For each approach, clustering coefficient, path length and rich-club coefficient were used to classify participants. Small-worldness was excluded as a composite of clustering and path length (Rubinov & Sporns, 2010). To ensure robust performance estimates, we employed 10-fold stratified cross-validation repeated 50 times, with stratification to maintain the original age group proportions in each fold. Performance was evaluated using classification accuracy.

#### 2.6.4 Network Topology-Cognition Associations

To investigate the relationship between network topology and and cognitive performance, we computed Pearson correlations between each graph metric and the cognitive composite, FDR-corrected across approaches. The stop signal and response inhibition reaction time measures were first multiplied by -1 so that higher scores represent better performance.

To evaluate the combined predictive power of graph metrics for cognition, ridge regression models were fitted with clustering coefficient, path length, and rich-club as predictors. Models were fit separately for each approach with 10-fold cross-validation to select the optimal regularization parameter (λ). Bootstrap resampling (n = 1,000) was used to estimate 95% confidence intervals for R² values. To isolate the contribution of network metrics beyond age, models were additionally run with age as a covariate.

To quantify whether graph metrics contributed cognitive-relevant information beyond age, we performed an F-change test comparing a full model (age + clustering + path length + rich-club) against a reduced model (age only). We also computed a bootstrap 95% confidence interval (n = 1,000) on the same quantity (ΔR²) to obtain a non-parametric estimate of reliability. Confidence intervals excluding zero indicate a reliable effect of topology beyond age.

To identify which graph metrics carried the cognitive signal, we also bootstrapped the standardised ridge coefficients for each approach and metric (n = 1,000). This was done separately for two models to quantify how much of each metric’s apparent cognitive relevance was shared with age: (i) a model with graph metrics only, and (ii) a model with graph metrics plus age. For each model, the β coefficient and 95% bootstrap CI were computed per predictor. Metrics whose 95% CI excluded zero were considered reliable individual predictors.

#### 2.6.5 Network Topology-Structural Connectivity Associations

For structural connectivity analyses, matrices were thresholded to 15% sparsity to match the metabolic connectivity and only edges present in >50% of subjects were retained in the group matrix (de Reus & van den Heuvel, 2013). For each connectivity approach, DICE coefficients were computed against the group-average SC connectome to quantify edge overlap.

##### Hub Analysis

SC hubs were defined as the top 25% of regions by weighted degree (van den Heuvel & Sporns, 2011; Wang et al., 2018). *Hub overlap* was calculated as the percentage of SC hubs that were also in the top 25% by degree in each connectivity approach. A *hub to non-hub ratio* was also computed as the mean degree of SC hub nodes divided by the non-hub nodes, providing a cross-modality index of whether SC hubs preserve their hub status (ratio > 1) or show an inverse relationship (ratio < 1) in each connectivity approach.

##### SC Prediction Analysis

To model the relationship between structural and metabolic regional edge strength in the group connectomes, we employed ridge regression with SC edge strength as the predictor and metabolic edge strength the outcome. This method applies L2 regularisation, which is particularly suitable as structural connectivity edge weights exhibit a heavy-tailed distribution. The shrinkage penalty stabilises coefficient estimates, mitigating the influence of extreme values. The optimal regularisation parameter (λ) was selected via cross-validation. Model performance was quantified as R² and beta coefficients were used to assess the direction and strength of SC-connectivity relationships.

#### 2.6.6 Network Topology-Gene Expression Associations

Lastly, we investigated the genetic basis of the connectivity approaches. For each approach, we computed region-level nodal degree centrality, defined as the number of edges connecting a region to other regions. We selected nodal degree for two reasons. First, it directly supports our aim of testing whether regional gene expression is associated with regional connectedness, while limiting the statistical family to six tests (one per approach). Second, degree centrality is defined for all 100 regions in every approach, whereas other metrics, such as nodal average shortest path length, were variably undefined for disconnected regions across approaches, which would confound the across-approach comparisons.

For each approach, Partial Least Squares Correlation (PLSC) was performed between the degree centrality vector and the gene expression matrix (100 regions × 94 genes). Significance was assessed using permutation tests with n = 1,000 permutations. A naive permutation test shuffled the region labels of the metric vector, breaking both spatial and network structure. A spatially-constrained spin permutation test was also used (Alexander-Bloch et al., 2018), which rotated the 100 Schaefer cortical region centroids on the spherical surface and reassigned region labels by nearest-neighbour matching, preserving the spatial autocorrelation inherent in cortical maps. Bonferroni correction was applied across the six approaches. Bootstrap resampling (n = 1,000) was also performed to estimate 95% confidence intervals for each gene’s loading. Genes with a confidence interval not crossing zero were considered statistically reliable.

## 3. Results

### 3.1 Characterising Network Topology of Metabolic Connectivity Approaches

We begin by describing the topological characteristics of each connectivity approach, quantifying their spatial similarity using DICE coefficients and comparing graph metrics across approaches.

The group-average connectomes for each connectivity approach are shown in Figure 2A. The two covariance matrices (Across-Subject and LOO) showed almost perfect alignment (DICE = 0.99), reflecting their shared covariance structure (Figure 2B left). For the other approaches, edge overlap was low-to-moderate (DICE 0.04-0.43), indicating that each connectivity method produced a relatively distinct spatial pattern of connections. fMRI Connectivity showed the highest similarity to metabolic approaches (mean DICE vs all other approaches = 0.34), while Kinetic C2 showed the lowest (mean DICE = 0.10), suggesting that metabolic kinetics are the most spatially distinct from other connectivity measures.

**Figure 2.**
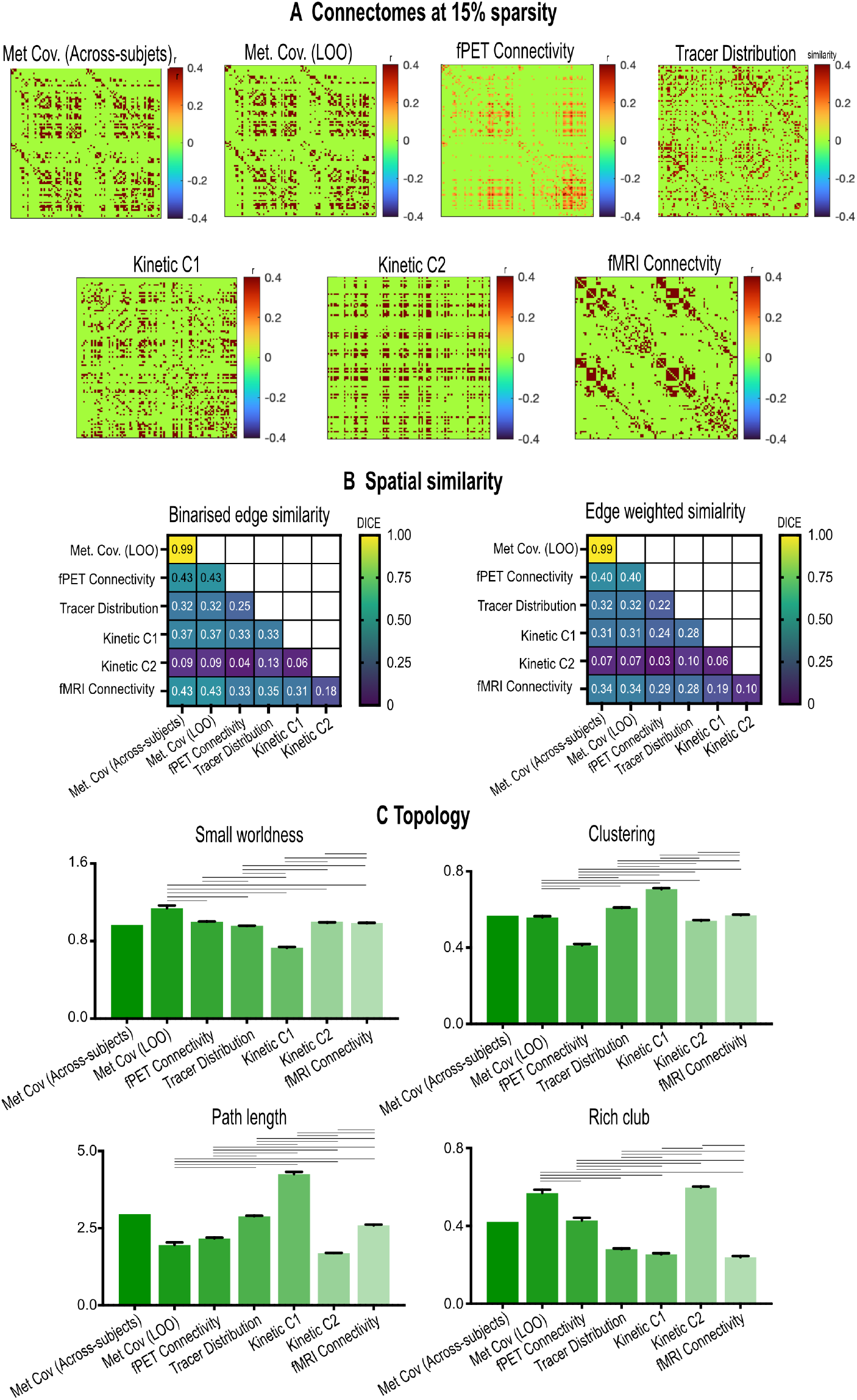
Characteristics of connectomes derived from connectivity methods. (A) Connectomes at 15% sparsity for each connectivity approach, including group-average metabolic covariance (Metabolic Cov. Group) as a reference. Edge weights represent Pearson r for Metabolic Cov (Across-Subject and LOO); Fisher-z(r) for fPET, fMRI and Kinetic C1 and C2; and a 0-1 distance-similarity index for Tracer Distribution. (B) Spatial similarity between connectomes assessed using binary DICE coefficients and edge-weighted coefficients, (Fuzzy DICE, which preserve edge weight information). (C) Network topology characterised by four graph metrics: small-worldness, clustering coefficient, characteristic path length, and rich-club coefficient. For each metric, repeated-measures ANOVA revealed significant differences across approaches (all p < 0.05). Horizontal lines above the bars indicate significant pairwise differences based on Bonferroni-corrected post-hoc comparisons (p < 0.05). Metabolic Cov. (Group) is shown for visual reference only and was not included in the ANOVA.

We additionally computed fuzzy DICE coefficients, which preserve connectivity edge strength, providing a measure of continuous similarity (Figure 2B right). With the exception of the two Metabolic Covariance approaches, Fuzzy DICE revealed low-to-moderate similarity across approaches (range: 0.03-0.40). Kinetic C2 again showed the lowest similarity to all other approaches (DICE = 0.03-0.10). This pattern confirms that the spatial distinctiveness of each connectivity approach is not simply an artefact of binarisation, but reflects true differences in the connectivity strength patterns across the methods.

We next explored the connectome from each approach using the graph metrics. The repeated-measures ANOVA for each graph metric (F(5,395) > 112.0, all p < 0.001) and Bonferroni-corrected post-hoc tests were significant, indicated that the connectivity approaches capture distinct network topologies (Figure 2C). fPET connectivity and Kinetic C2 networks both exhibited small-worldness with values at or above the canonical threshold of 1.0 (σ = 1.00 for both), indicating an optimal balance between local specialisation and global integration (also see Supplement 7). Metabolic Covariance (LOO) demonstrated the strongest small-worldnesss (σ = 1.13), driven by high clustering (CC = 0.56) and relatively short path lengths (L = 1.96), suggesting particularly efficient network architecture. Interestingly, the Metabolic Covariance (Across-Subject) network (σ = 0.97) fell slightly below the threshold, highlighting that the LOO method enhances small-world architecture while not substantially altering the overall topological profile. In contrast, Tracer Distribution (σ = 0.96) and fMRI Connectivity (σ = 0.99) fell just below the small-world threshold of 1.0. Most notably, Kinetic C1 (σ = 0.73) lacked small-world characteristics.

Collectively, these findings demonstrate that while most connectivity approaches produce near-optimal or optimal small-world topologies, the degree of local segregation versus global integration varies substantially across methods. These topological differences suggest that the choice of connectivity method fundamentally alters the observed organisational principles of brain networks, with important implications for comparing results across studies that employ different approaches. This methodological sensitivity is particularly critical given that the topological signatures we observed would lead to fundamentally different conclusions about the organisational principles of the same brain network, from highly clustered but weakly integrated (Kinetic C1) to globally efficient and hub-dominated (Metabolic Covariance (LOO)). Researchers should therefore exercise caution when directly comparing network properties across studies that use different connectivity measures, and consider whether observed differences reflect genuine biological variation or methodological choice.

### 3.2 Age Group Differences in Network Topology of Metabolic Connectivity Approaches

Healthy ageing is associated with distinct patterns of alterations to cerebral glucose metabolic (Deery et al., 2023a), functional connectivity (Deery et al., 2023b) and metabolic connectivity (Deery et al., 2025b). As such, determining each method’s sensitivity to age-related change is a useful model to test its measurement validity. Hence, we compared younger (N=36) and older (N=44) adults across graph metrics using independent samples *t*-tests (Figure 3A) and quantified the utility of these metrics for classifying age group membership using discriminant analysis (Figure 3B).

**Figure 3.**
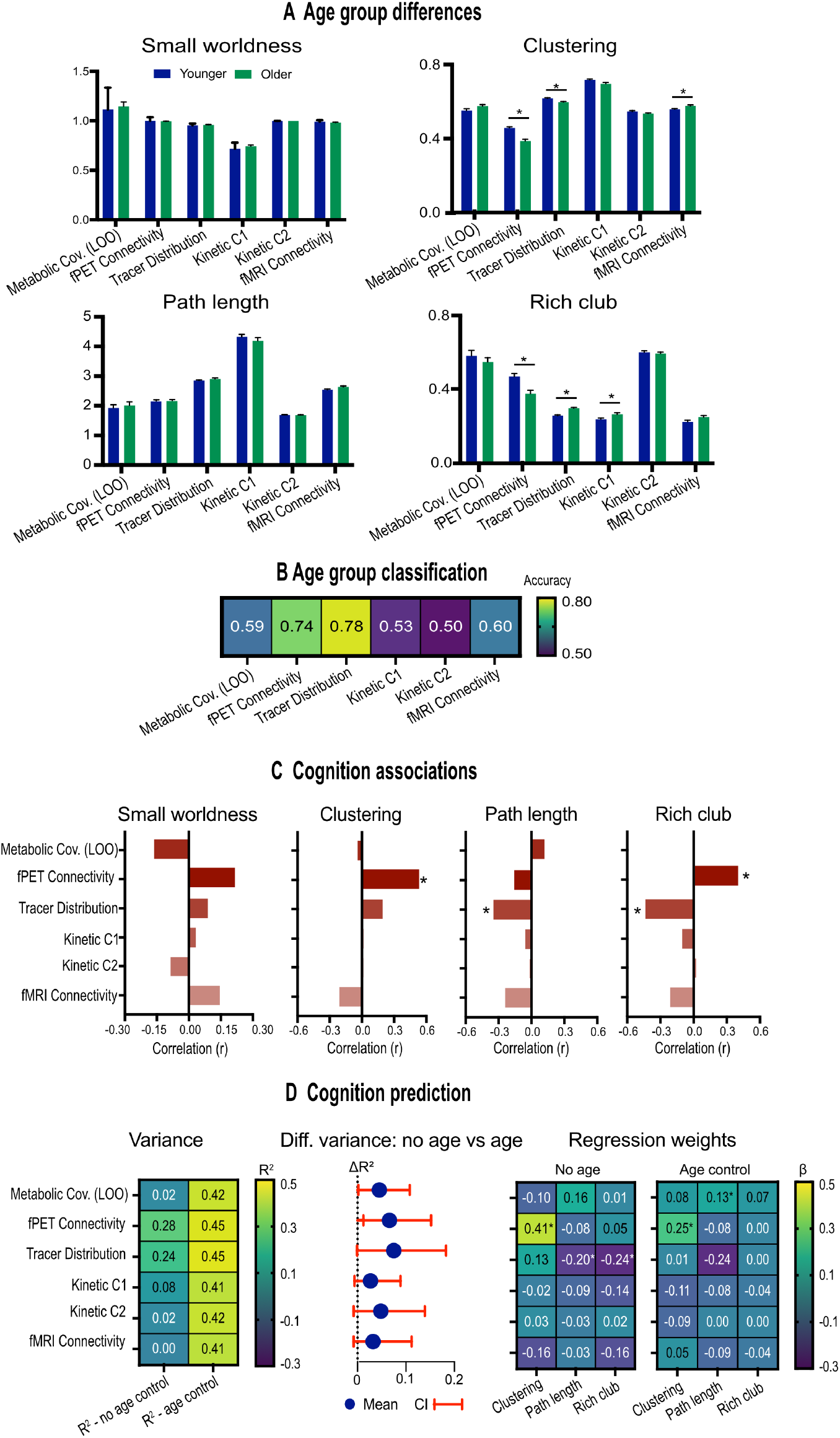
Age group differences and cognition associations across connectivity approaches. (A) Age group differences in graph metrics. Bars represent mean values with standard error of the mean (SEM) for each metric. Asterisks (*) indicate significant age differences (p-FDR < 0.05). (B) Age-group classification accuracy from discriminant analysis based on clustering coefficient, path length, and rich-club. (C) Pearson correlations between graph metrics and the PCA-derived cognitive composite score (PC1, explaining 39.6% of variance). Bars show correlation coefficients *(r);* asterisks (*) indicate significant correlations (p-FDR < 0.05). (D) Ridge regression results predicting cognition from graph metrics. Left panel shows variance explained (R²) for models without age and with age as a covariate; middle panel is mean and confidence intervals for difference in R^2^ between no age and age models; right panel is coefficients for the graph metrics, with asterisk (*) indicating non-overlapping bootstrap 95% confidence intervals from zero (i.e., significant predictive performance).

Tracer Distribution emerged as the strongest classifier (accuracy = 78%), consistent with large age group differences for rich-club (Cohen’s d = -1.45, p < 0.001) and clustering (d = 0.86, p < 0.001). The opposing direction of these effects (higher rich-club but lower clustering in older adults) suggests a fundamental reorganisation of voxel-level distribution patterns with ageing, shifting from the strongly clustered, segregated topology observed descriptively in the whole sample (C = 0.62, L = 2.88) toward a more hub-dominated configuration in older people.

fPET connectivity showed the second-highest classification accuracy (74%), with age group differences in clustering (d = 1.23, p-FDR < 0.001) and rich-club (d = 0.85, p < 0.001), both showing lower values in older adults. These strong age group differences are notable given that fPET Connectivity showed moderate local clustering in the whole sample (C = 0.42), suggesting that even modest clustering in dynamic metabolic networks is highly vulnerable to age-related decline and provides robust discriminative power.

fMRI connectivity achieved moderate classification accuracy (60%), consistent with its significant age-related differences in clustering (d = -0.63, p-FDR = 0.007) and path length (d = -0.52, p-FDR = 0.024). These findings align with the fMRI literature on age-related dedifferentiation and loss of network efficiency in older adults (Deery et al., 2023b). Kinetic C1 also showed significant but weaker age group differences, with rich-club higher in older adults (d = -0.53, p-FDR = 0.021), yet achieved only classification accuracy modestly above chance (53%), suggesting that the age-related shift toward hub-dominated configuration is not sufficient for reliable individual-level classification.

Metabolic Covariance (LOO) and Kinetic C2 showed no significant age group differences in the graph metrics. The classification of Kinetic C2 (50%) was essentially at chance level, confirming that this approach captures little age-related information despite showing short path lengths and rich-club characteristics in the whole sample. Metabolic Covariance (LOO) achieved moderate accuracy (59%) despite no significant univariate age effects for the graph metrics. This moderate multivariate discriminative power suggests that network properties of Metabolic Covariance (LOO) contain subtle age-related information when considered in combination rather than as individual features.

### 3.3 Cognition-Network Topology Associations

Conceptually, functional and metabolic connectivity are expected to show associations with cognition (Jamadar, 2026) although effect sizes and reliability is a matter of debate (Chopra et al., 2024). To examine the relationship between graph metrics and cognitive performance, we computed Pearson correlations with a PCA-derived composite of cognition (see Methods). Across approaches, significant associations with cognition were found for fPET Connectivity and Tracer Distribution only and revealed distinct and opposing patterns.

For fPET Connectivity, higher clustering was associated with better cognition (r = 0.53, p-FDR < 0.001), the largest effect observed across all approaches and metrics (Figure 3C). Higher rich-club was also associated with better cognition (r = 0.39, p-FDR = 0.001). These findings indicate that higher local clustering and hub organisation in fPET Connectivity networks are associated with better cognitive performance. Like with the age differences, these cognition effects also suggest that even modest absolute levels of local clustering observed in fPET networks have predictive utility for cognition.

In contrast, Tracer Distribution showed significant negative associations with cognition for rich-club and path length (r = -0.44 and r = -0.35, both p-FDR < 0.01), indicating that higher hub organisation and shorter path lengths (more integration) are associated with better cognition. This pattern is notable given that the tracer distribution network in the whole sample was characterised by high clustering and long path lengths, suggesting that deviation from this segregated topology toward greater integration is cognitively beneficial, while increased hub organisation is maladaptive. No other approach showed significant cognitive associations after FDR correction.

For the ridge regression analyses, fPET Connectivity and Tracer Distribution were the only approaches that predicted cognition above chance (Figure 3D left), explaining approximately 28% and 24% of variance in cognitive performance, respectively. Clustering in fPET was the strongest positive predictor (β = 0.41), while Tracer Distribution (path length, β = -0.20; rich-club, β = -0.24) were the strongest negative predictors.

To determine whether the cognitive-relevant topology of each approach was distinct from age-related variance, we compared a full model (age + clustering + path length + rich-club) with a reduced model (age only) using two complementary approaches: a parametric *F*-change test on the ΔR² and a bootstrap 95% confidence interval. Age alone explained ∼40% of cognitive variance across approaches. Adding topological metrics improved prediction by 5-6% for fPET Connectivity and Tracer Distribution, and by 1-3% for the remaining approaches (Figure 3D middle). fPET Connectivity (F(3, 75) = 2.29, p = 0.085) and Tracer Distribution (F(3, 75) = 2.63, p = 0.057) both showed trend-level effects. Bootstrap 95% CIs on ΔR² excluded zero for fPET Connectivity (point estimate 0.051, 95% CI (0.012, 0.152)) and Metabolic Covariance (LOO) (0.026, (0.002, 0.108)).

The two tests therefore agreed on fPET Connectivity, both pointing toward an effect of topology beyond age. However, the tests disagreed on Metabolic Covariance (LOO) (bootstrap CI excluding zero, but F-change p = 0.346) and Tracer Distribution (F-change trend, but bootstrap CI including zero). The Metabolic Covariance (LOO) result should be interpreted with caution because the ridge regularisation parameter was unstable across cross-validation folds (λ capped to the search boundary), and the LOO-derived topology metrics showed anomalous between-subject variance (path length SD = 0.73, three- to four-fold larger than all other approaches), suggesting that the apparent effect may be an artefact of the LOO procedure rather than a genuine cognitive association. We therefore treat fPET Connectivity as the strongest evidence for topology-specific cognitive prediction beyond age, with Tracer Distribution showing concordant but trend-level evidence.

To identify which individual graph metrics most contributed to the cognitive effect, and whether those associations were independent of age, we bootstrapped the standardised ridge coefficients for each approach in both the without-age and with-age models (Figure 3D right). In the without-age model, four of the 18 tested coefficients had 95% CIs that excluded zero, including clustering in fPET Connectivity, clustering in Tracer Distribution, path length in Tracer Distribution, and rich-club in Tracer Distribution. However, when age was included in the model, only clustering in fPET Connectivity and path length in Tracer Distribution remained significant.

Collectively, these findings demonstrate that while age is a strong predictor of cognition across all approaches, only fPET Connectivity provides robust topology-specific information about cognition beyond age. Furthermore specific graph metrics provide robust associations with cognition, namely, fPET Connectivity’s nodal clustering (local segregation) and Tracer Distribution’s path length (global integration). The remaining approaches, and the remaining metrics, primarily reflect age-related variance, a pattern that suggests both the intrinsic constraints of the covariance-based approach and methodological issues arising from the LOO estimation procedure.

### 3.4 Structural Connectivity and Metabolic Connectivity Approaches

Metabolic connectivity, as a form of functional connectivity, emerges from a spatially constrained neuroanatomical scaffold of neurons, axons, fibre bundles and tracts (Suarez et al., 2020; van den Heuvel et al., 2015). As such, it seems reasonable to expect some level of association between structural and metabolic connectivity (Jamadar, 2026; Jamadar et al., 2025). Indeed, there exists a non-trivial association between structural and fMRI-derived measures of functional connectivity, but the two are not fully concordant (Suarez et al., 2020), with associations (r) at the individual subject level between 0.02-0.25 (Straathof et al., 2019). These low associations reflects the fact that structure-function relationships emerge through both lower-order (two nodes may interact directly via a shared connection) and higher order (two nodes may interact indirectly via common neighbours) interactions. As such, one would expect that structural and metabolic connectivity to be related, but not perfectly so. We test these associations next.

#### 3.4.1 Edge Overlap and Strength Similarity

DICE coefficients revealed uniformly low-to-moderate edge overlap between SC and all connectivity approaches (range: 0.10-0.50) (Figure 4A). fMRI showed the highest overlap with SC (DICE = 0.50), followed by Metabolic Covariance (LOO; DICE = 0.40) and fPET Connectivity (DICE = 0.37). Tracer Distribution (DICE = 0.27) and Kinetic C1 (DICE = 0.32) showed lower overlap, while Kinetic C2 showed the weakest spatial alignment with SC (DICE = 0.10). This pattern suggests that fPET Connectivity and Metabolic Covariance (LOO) most closely mirror structural architecture, while glucose kinetics are largely independent of structural connectivity patterns.

**Figure 4.**
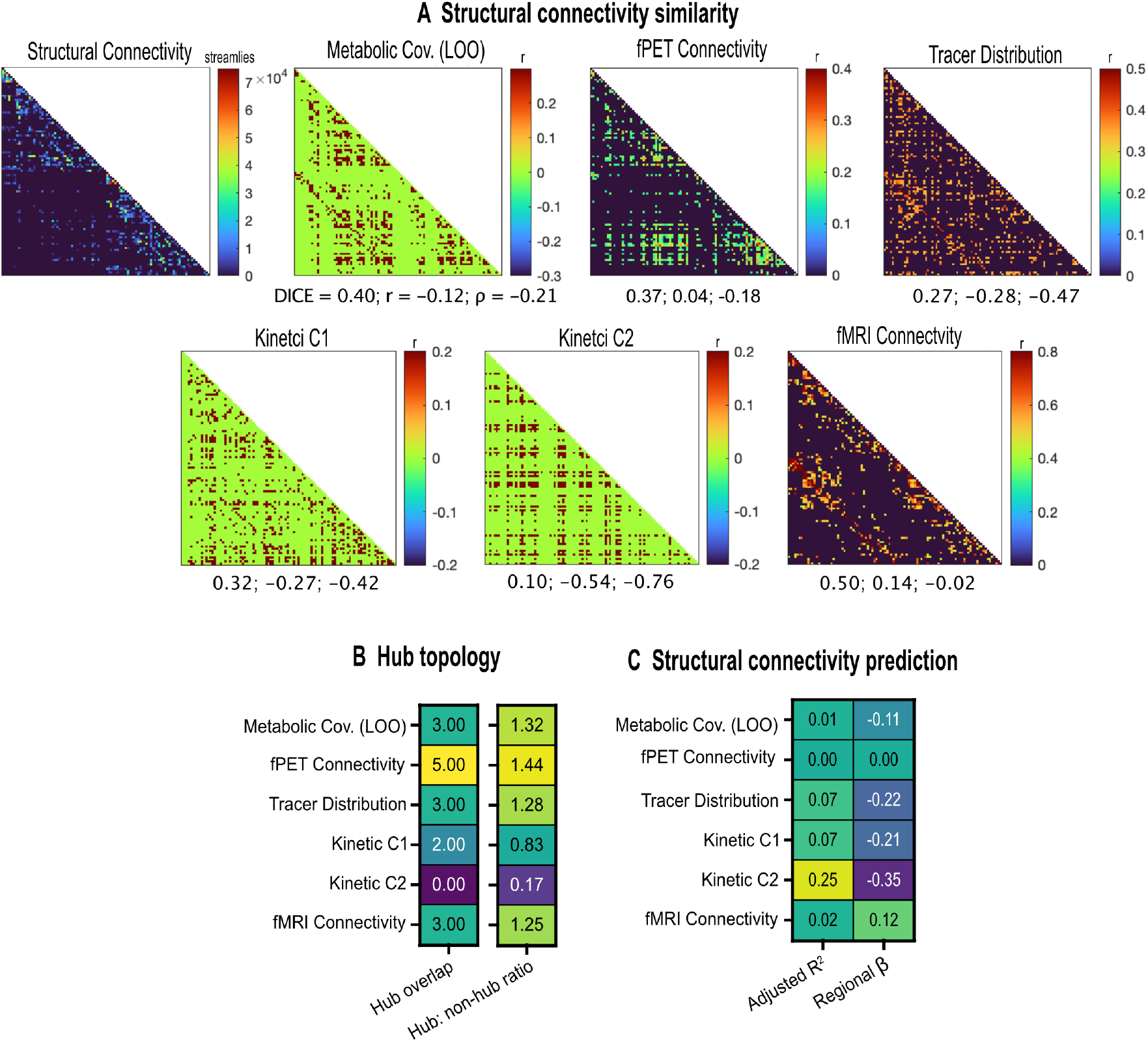
Structural connectivity associations across connectivity approaches. (A) Regional structural connectome (number of streamlines between regions) and similarity with connectome in each approach. Beneath each connectome: spatial similarity (DICE), edge-wise Pearson correlation (r), and Spearman’s rank correlation (ρ). (B) Hub topology similarity between SC and each connectivity approach. Left column shows the number of hubs in SC matrix (top 25% of regions by weighted degree) shared with each metabolic approach; right column shows the hub: non-hub degree ratio in the connectivity approaches (values > 1 indicate SC hubs retain hub status in the metabolic approach; values < 1 indicate anti-hub organisation). (C) Ridge regression results predicting connectivity in each connectivity approach from regional structural connectivity. Left column is variance explained (R²) with significance asterisk (*) from permutation testing (n=1,000). fPET Connectivity was the only approach that did not significantly predict structural connectivity (p = 0.229). Right column is standardised beta coefficients (β) from the ridge regression models. All approaches showed negative associations, except fMRI, which showed a positive association.

We computed Pearson correlations between the SC and connectivity network edges. Given the heavy tailed distribution of SC, non-parametric Spearman correlations were also computed. Spearman correlations were consistently more negative than Pearson correlations across all approaches (Figure 4A). For fMRI Connectivity, the positive Pearson correlation (r = 0.14) became near-zero (ρ = -0.02), while for all other approaches, Pearson correlations became more negative in the Spearman rank-order analysis: fPET Connectivity (r = 0.04 vs ρ = -0.18), Metabolic Covariance (LOO) (r = -0.12 vs ρ = -0.21), Tracer Distribution (r = -0.28 vs ρ = -0.47), Kinetic C1 (r = -0.27 vs ρ = -0.42), and most strikingly, Kinetic C2 (r = -0.54 vs ρ = -0.76). These results indicate that the relationships between SC and metabolic connectivity are not simply linear; rather, the strongest effects are driven by rank-order patterns more than raw edge-weight correspondence. The consistently negative Spearman correlations suggest that, across approaches, edges with higher SC strength tend to have lower metabolic connectivity strength, and vice versa.

#### 3.4.2 Hub Analysis

SC hubs were defined as the top 25% of regions by weighted degree (van den Heuvel & Sporns, 2011; Wang et al., 2018). These hubs were distributed across multiple large-scale networks, including visual, somatomotor, dorsal attention, default mode, salience, and control networks. Hub overlap between SC and metabolic approaches was modest (Figure 4B left): fPET Connectivity showed the highest overlap (20%, 5 of 25 SC hubs), followed by Metabolic Covariance (LOO), Tracer Distribution, and fMRI Connectivity (each 12%, 3 hubs). Kinetic C1 showed lower overlap (8%, 2 hubs), while Kinetic C2 showed no overlap with SC hubs (0%).

We also computed a *hub to non-hub ratio* by comparing the degree of SC hubs and non-hubs. While hub overlap reveals discrete topological patterns of shared regional hubs, the hub to non-hub ratio quantifies the nature of those hub connections. fPET Connectivity (1.44), Metabolic Covariance (LOO) (1.32), Tracer Distribution (1.28) and fMRI Connectivity (1.25) all showed ratios > 1.0, indicating that SC hubs also have higher metabolic connectivity degree on average than non-hubs (Figure 4B left). Together, these results suggest that while SC hub status does not entirely predict metabolic hub status, SC hub regions are generally more metabolically connected than non-hubs. In contrast, Kinetic C1 showed a ratio below 1 (0.83), suggesting that SC hubs have lower degree in kinetic dynamics than non-hubs or a mild anti-hub pattern. Strikingly, Kinetic C2 showed a ratio near zero (0.17), indicating that SC hubs are strongly anti-hubs in Kinetic C2 connectivity. In other words, regions that are structural hubs have much lower connectivity in Kinetic C2 compared to non-hubs.

#### 3.4.3 SC Prediction of Metabolic Connectivity

In ridge regression, Kinetic C2 Connectivity showed the strongest prediction of SC connectivity (R² = 0.25, β = -0.35), followed by Tracer Distribution (R² = 0.07, β = -0.22) and Kinetic C1 (R² = 0.07, β = -0.21) (Figure 4C). The negative beta coefficients for Kinetic C2 confirm that the relationship between SC and Kinetic C2 is inverse: stronger structural connections predict weaker C2 TAC similarity. fMRI (R² = 0.02, β = 0.12) and Metabolic Covariance (LOO) (R² = 0.01, β = -0.11) showed weak prediction of SC connectivity, while fPET Connectivity showed no predictive power (R² ≈ 0, β ≈ 0).

The strong predictive performance of Kinetic C2 for SC, despite its poor alignment with SC (lowest DICE, negative edge correlations, 0% hub overlap), represents a key finding. This apparent paradox may resolve if it is accepted that independence from structural constraints does not imply randomness. Rather, C2 TAC distance captures metabolic processes that are systematically, albeit inversely, related to the brain’s structural architecture. This is consistent with the interpretation that the similarity of glucose kinetic TACs operate according to a different organisational principle than other approaches. That principle is not spatially constrained by structural connectivity but is still systematically (and inversely) related to it.

### 3.5 Mapping of Metabolic Gene Expression and Metabolic Connectivity Topology

Given the known physiology underlying FDG-PET (Hahn, 2025; Yakushev et al., 2017), one would predict that metabolic connectivity would be associated with regional expression of genes associated with metabolic processes. This has recently been shown for fPET metabolic connectivity gradients (Deery, Moran, et al., 2026a). To test these associations across the connectivity approaches, we performed Partial Least Squares Correlation (PLSC) of the regional expression of 94 glucose-metabolism genes and nodal degree centrality. Significant associations were found for all six approaches (spin-corrected p < 0.001; Figure 5A left), with two clearly opposing directions of association. Positive associations were found for Metabolic Covariance (LOO) (r = 0.65), Tracer Distribution (r = 0.51), and fPET Connectivity (r = 0.50), indicating that regions with greater connectivity also expressed higher levels of the glucose-metabolism gene signature. Negative associations were found for Kinetic C1 (r = -0.57), fMRI Connectivity (r = -0.53), and Kinetic C2 (r = -0.53), indicating the opposite.

**Figure 5.**
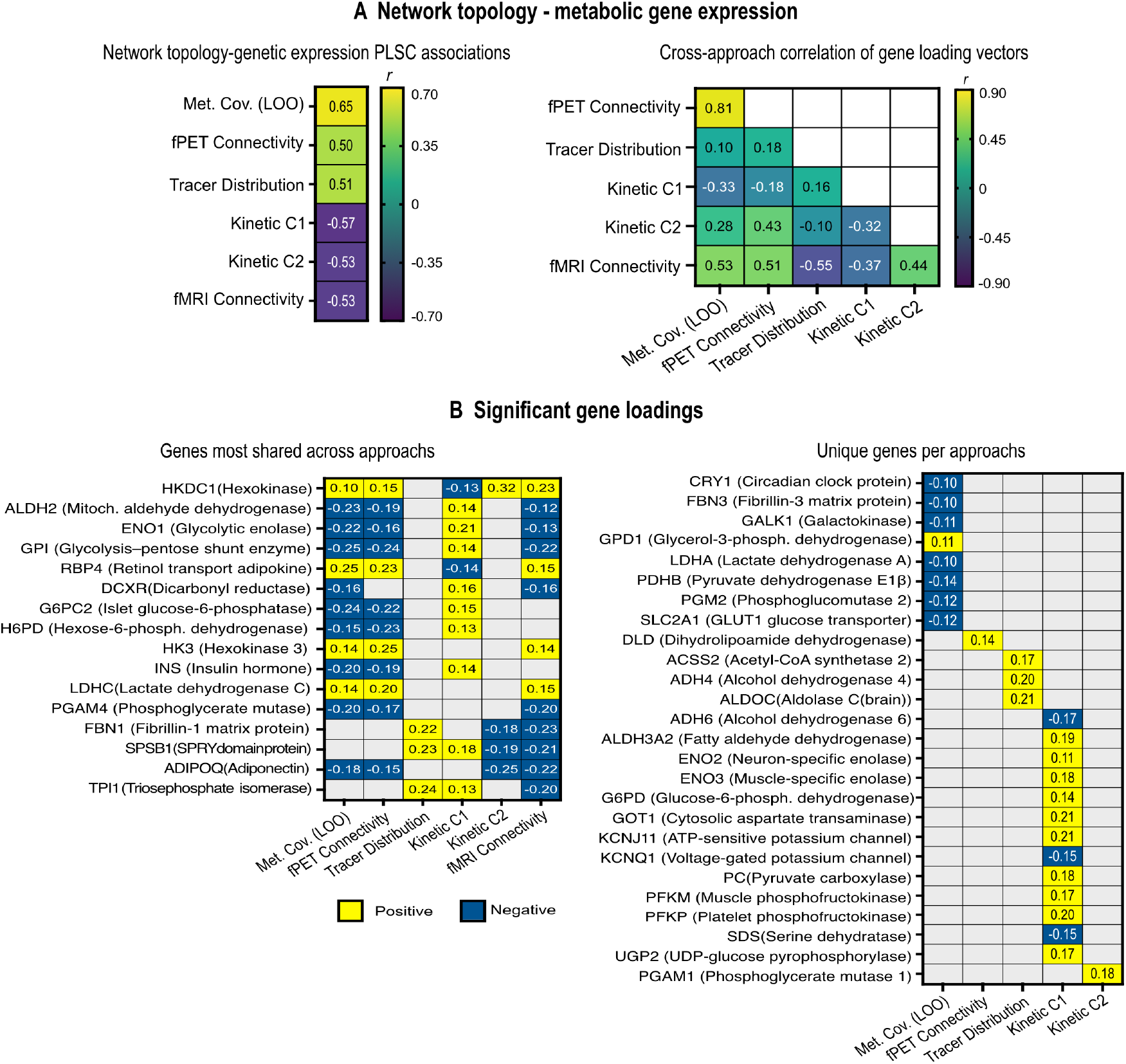
Gene expression association in the connectomes of the connectivity approaches. (A) Partial Least Squares Correlation (PLSC) between network topology (nodal degree) and 94 glucose metabolism genes in each approach. All approaches were significant (permutation tests, p < 0.05). (B) Cross-approach correlation matrix of gene loading vectors. Values are Pearson correlations (r) between the 94-element loading vectors of each pair of approaches; positive values indicate convergent gene signatures; negative values indicate opposing signatures. (C) Genes most shared across approaches. Rows are genes significant in three or more approaches; cells show the loading value per approach, with colour indicating sign and blanks indicating non-significant loadings. (D) Approach-specific gene signatures. Rows are genes significant in only one approach; cells show the loading value, coloured by sign. Full gene results and enrichment analyses are available in Supplement 8.

Bootstrap resampling identified reliable gene associations of varying breadth: 33 genes for Metabolic Covariance (LOO); 28 for Kinetic C1; 18 each for fPET Connectivity and fMRI Connectivity; 13 for Kinetic C2; and seven for Tracer Distribution. Breadth did not perfectly track overall association strength, with Tracer Distribution showing a strong regional correspondence (r = 0.51) with only seven significant genes. Sorting the reliably loaded genes by the number of approaches in which they appeared revealed a shared core of 16 genes significant in three or more approaches (Figure 5B left). Functionally, these genes span glucose handling, with hexokinases that initiate glycolysis; glycolytic enzymes; glucose-6-phosphatase that opposes glycolysis and gates insulin secretion; dehydrogenases that manage metabolic by-products; and the hormone insulin (INS). Collectively this gene set constitutes a coherent glucose-sensing and insulin-signalling network rather than a random subset of metabolic genes.

Comparison of gene loading vectors revealed an aligned with two groups of approaches (Figure 5A right). For the first group, Metabolic Covariance (LOO) and fPET Connectivity showed the strongest convergence (r = 0.81), reflecting a shared glucose-metabolism signature. Both loaded with RBP4 positively and GPI, G6PC2, ALDH2, and INS negatively (Figure 5B, left). For the second group (Kinetic C1 and Kinetic C2), these shared core genes had the opposite sign. Functionally, RBP4 and RBP4 are a retinol transporter and an insulin-resistance adipokine; GPI sits at the crossroads of glycolysis and the pentose phosphate pathway and has neuroprotective roles; G6PC2 is an islet-specific enzyme that acts as a brake on insulin release; and ALDH2 is a mitochondrial aldehyde dehydrogenase whose deficiency has been associated with cognitive impairment. These results reflect a shared glucose-metabolism signature across groups of approaches but associations in the opposing direction.

fMRI Connectivity was the sole approach whose loading vector bridged the clusters. It correlated positively with Metabolic Covariance (LOO) (r = 0.53) and fPET (r = 0.51), indicating a shared gene signature, yet fMRI’s degree association was negative like the kinetic cluster. This dissociation places fMRI at the bridge between the two clusters. The two kinetic approaches showed weak loading correlation with each other (r = 0.18). They shared only HKDC1 and SPSB1, which loaded in opposite directions. Tracer Distribution showed the most divergent profile. Despite a positive degree association similar in magnitude to fPET Connectivity (r = 0.51), its loading vector was uncorrelated with the metabolic cluster (r = 0.10) and strongly anti-correlated with fMRI (r = -0.55).

Approach-specific signatures further differentiated the approaches (Figure 5B right). Kinetic C1 carried the largest unique signature (13 genes), dominated by glycolytic flux enzymes, reinforcing its distinct emphasis on glycolytic metabolism. Kinetic C2, despite sharing the kinetic modelling framework with C1, had a much smaller and largely distinct signature of one unique gene (PGAM1). Metabolic Covariance (LOO) also carried a substantial unique signature (8 genes), including a broad set of genes consistent with insulin and metabolism. Tracer Distribution’s significant genes are largely structural/transport-related rather than insulin-glucose axis genes, suggesting this approach captures a distinct biological feature. fMRI Connectivity had no uniquely loaded genes, which is consistent with its intermediate loading correlations with both clusters.

Pathway enrichment analysis (Zhou et al., 2019) of the reliably loaded genes was undertaken to provide additional descriptive information across approaches (Supplementary Table S5). The top enriched term for every approach was a glucose-metabolism pathway, including glycolysis/gluconeogenesis, glucose metabolic process, or carbohydrate metabolic process. Beyond this shared core, approaches diverged in breadth and specificity. Kinetic C1 showed the broadest and most diverse metabolic enrichment, including glycerol metabolism, galactose metabolism, NAD+ metabolism, and the Cori cycle. Metabolic Covariance (LOO) uniquely enriched for AMPK and glucagon signalling, extending its association to whole-body glucose regulation. fPET Connectivity showed the highest percentage of genes in glycolysis (68%) and uniquely enriched for HIF-1 signalling (a key glycolytic regulator) and genes implicated in Type II diabetes. Kinetic C2’s smaller signature was dominated by glucose homeostasis and pyruvate metabolism, consistent with its oxidative/gluconeogenic profile. fMRI Connectivity showed strong glucose-metabolism enrichment (74% glucose metabolic process) despite its negative degree association, reinforcing that its gene signature resembles the metabolic approaches but with inverted topology-gene correspondence.

Collectively, these findings indicate that the direction of the connectivity-gene association is a primary feature differentiating the connectivity approaches. All six approaches implicated a shared core glycolytic and insulin machinery, but the sign of the association with regional connectedness split into two groups: the metabolic approaches (Metabolic Covariance (LOO) and fPET Connectivity), plus Tracer Distribution, showed positive associations, whereas fMRI Connectivity and the kinetic approaches (C1 and C2) showed negative associations. The unique gene signatures indicated that Metabolic Covariance (LOO) and fPET Connectivity emphasise the insulin-resistance axis, kinetic methods emphasise glycolytic flux, Tracer Distribution captures structural/transport-related genes, and fMRI draws entirely on genes shared with other approaches.

### 3.6 Sensitivity Analysis: Results at 25% Network Sparsity

To assess the robustness of our findings, we repeated the main analyses at a higher network sparsity threshold of 25% (see Supplement 9). The key topological, cognitive, and structural findings were concordant across sparsity levels. The genetic associations were preserved for Metabolic Covariance (LOO), fPET Connectivity, Kinetic C1 and Kinetic C2. Tracer Distribution and fMRI Connectivity showed sparsity-dependent associations, with their network degree-gene direction and placement within the two-cluster structure reversing between thresholds. Taken together, these findings support the main conclusions that fPET Connectivity is the approach most tightly coupled to cognition and to age-related decline. It shares spatial overlap with structural connectivity and maps to metabolic gene expression. The Kinetic C1 and C2 approaches show minimal relationship with cognition and in some cases opposite structural associations, while fMRI Connectivity provides complementary insights into structural and genetic associations with network topology. Despite strong topological features and gene associations, Metabolic Covariance (LOO) shows comparatively weaker brain-behaviour relationships. Tracer Distribution and fMRI Connectivity approaches also appear uniquely sensitivity to the choice of edges retained at threshold when mapped to metabolic gene expression.

## 4. Discussion

The results of this study show that the choice of approach for calculating metabolic network connectivity from FDG-PET data is not merely a technical decision. It fundamentally determines which brain-behaviour and brain-physiology relationships are observable. Our comparison of six metabolic connectivity approaches revealed that the methods yield divergent, and in some cases opposite, conclusions about the neural basis of cognition and ageing and the neurophysiological underpinnings of brain network organisation. By systematically evaluating network topology, cognitive associations, age sensitivity, structural and genetic alignment, we demonstrate that the optimal approach depends on the research question, with fPET Connectivity emerging as the most promising metabolic marker for cognition and ageing.

A key finding across our analyses is that greater topological strength alone does not always equal greater functional relevance. Metabolic Covariance (LOO) exhibited the highest small-worldness and rich-club organisation, indicating a highly clustered, hub-dominated and globally efficient network. However, despite these properties, it showed no univariate relationships with cognition nor age, and only moderate multivariate age group discrimination. These results confirm previous work indicating poor overlap between across-subject metabolic covariance networks and within-subject connectivity approaches (Jamadar, Ward, et al., 2021; Reed et al., 2023; Volpi et al., 2023) with our results showing that leave-one-out analysis is not sufficient to resolve true individual-level variance. These results caution against interpreting topological metric strength without considering the underlying biology and functional relevance of the connectivity measure. Our structural connectivity and gene expression analyses further suggests that different connectivity approaches are associated with distinct and in some instances opposing directions of associations. Together, these results support the need for considering multiple connectivity approaches to best interpret the nature of PET-derived networks.

fMRI-derived functional connectivity showed the strongest alignment with structural connectivity but largely weak cognitive associations, reinforcing that, like topological strength, structural alignment alone does not indicate cognitive relevance. The BOLD signal is an indirect marker of neuronal activity, comprising relative changes in cerebral blood flow, blood volume and oxygenation (Gauthier & Fan, 2019; Liu, 2013; Logothetis & Wandell, 2004). Our results are consistent with work indicating that neural activity elicits stronger responses in glucose metabolism than in oxygen consumption (Epp et al., 2025; Ogawa et al., 1990), highlighting the complementary value of PET-based methods beyond haemodynamic measures. Other fPET work has demonstrated robust increases in glucose metabolism across various cognitive tasks with high test-retest reliability (Hahn et al., 2016; Hahn et al., 2018; Hahn et al., 2024; Jamadar et al., 2019; Jamadar, Zhong, et al., 2021; Rischka et al., 2018). Together, this growing body of functional PET work supports the validity of dynamic metabolic connectivity measures, their complementary insights into network organisation, and their potential as biomarkers for cognitive ageing and disease.

The analyses of cognition with and without age as a covariate reveals that different topological metrics have different age-dependence. fPET Connectivity’s clustering coefficient and Tracer Distribution’s path length both survived age adjustment, indicating that these metrics capture cognition-relevant information that is not simply age-related. In contrast, Tracer Distribution’s clustering and rich club coefficients shrank substantially when age was included, revealing that their apparent cognitive relevance was largely accounted for by age-related variance in both cognition and topology. These findings suggest that topological metrics cannot be treated as interchangeable within a method. Rather, age-dependence of each specific metric must be characterised individually.

Our DICE coefficients for fMRI and the kinetic approaches were lower than those reported by Volpi and colleagues (Volpi et al., 2023). Possible sources of this discrepancy include differences in fMRI pre-processing pipelines, sparsity thresholds, atlas parcellations, and resting-state compared to naturalistic conditions during data acquisition. We also found Kinetic C2 presented an apparent contradiction: it showed the poorest spatial similarity with structural connectivity yet its network edge strength was the strongest predictor of structural connectivity strength, albeit reflecting the inverse spatial organisation. Kinetic C2’s positive unique association with PGAM1, a key enzyme in the glycolytic pathway, further supports the interpretation that C2 TAC similarity operates according to a different biological principle than structural or functional connectivity. Kinetic C2 also showed no significant cognitive associations, suggesting that this measure may capture unique biological processes but lacks cognitive relevance.

Collectively, these results suggest that the optimal connectivity approach depends on the research question. For cognition and ageing studies, our data indicates that fPET Connectivity is a robust choice as it shows large age effects and age-independent predictive power for cognitive performance. In contrast, Metabolic Covariance (LOO), despite being computationally straightforward, requiring only static SUVR images and standard correlation analyses, should be used with caution for brain-behaviour investigations. For multi-modal integration, fMRI functional connectivity offers strong structural alignment. Tracer Distribution is valuable for age classification and for single-scan designs where dynamic PET is not feasible, but requires careful interpretation of its negative age and mixed direction of cognitive associations. Kinetic approaches offer novel biological insights into glucose kinetics but similarity measures of regional TACs appear to lack predictive power for ageing and cognition compared to other approaches.

Although fPET Connectivity emerged as the most promising metabolic marker for cognition and ageing, the method has practical considerations. fPET Connectivity requires dynamic listmode PET acquisition with infusion or bolus+infusion radiotracer administration protocols, extended scan durations (typically 60+ minutes) for timeseries estimation and the analytic pipeline is more computationally demanding than single scan approaches. Despite these requirements, fPET Connectivity’s unique ability to capture temporal dynamics enables time-variant analyses that static covariance approaches cannot provide (Jamadar, Ward, et al., 2021). Recent work shows, for example, that information contained in the temporal dynamics of the signal is related to network efficiency and cognition (Deery, Liang, et al., 2026) and has utility for understanding how changes in glucose use support directed information flow in circuits supporting cognition and affective regulation (Deery, Moran, et al., 2026b). Analysing the temporal variability of the glucodynamic signal also enables the study of time-varying functional connectivity (sometimes called ‘chronnectomics’ (Calhoun et al., 2014) or ‘dynamic functional connectivity’) and metabolic microstates (Deery et al., 2025a). Approaches that use distance measures of kinetic TACs or tracer distribution are unable to resolve these time-variant neurobiological properties of cerebral glucose metabolism. Hence, fPET Connectivity is currently the only method for studies of temporal dynamics of brain metabolism (Jamadar, 2026; Jamadar et al., 2025).

The Tracer Distribution and Kinetic (C1 and C2) approaches reveal distinct and opposing connectivity patterns, suggesting they capture fundamentally different biological processes. The Tracer Distribution method, by analysing the full voxel intensity profile of regions, indexes regional heterogeneity and the ‘tails’ of the metabolic supply distribution. In contrast, the Kinetic approaches leverage the temporal trajectory of tracer uptake and clearance to reveal the dynamics of metabolic demand. The divergent patterns observed across these methods imply that they may serve as complementary biomarkers of distinct pathophysiological states. From a practical standpoint, Tracer Distribution offers the advantage of employing short-duration or single-image acquisitions (e.g., bolus radiotracer administration), as it requires only static PET data and obviates the need for dynamic scanning. Computationally, while it is more demanding than simple correlation-based metrics due to voxel-wise distribution extraction, energy distance calculations, and mutual information correction, it is simpler than full compartmental kinetic modelling. However, the negative associations with cognitive performance and positive associations with age observed in our data warrant careful interpretation. Higher “connectivity” in this context may reflect pathological tracer accumulation or metabolic dysregulation rather than functional integration, a notion consistent with the method’s original development for disease-state applications (Labarthe et al., 2026).

Kinetic approaches (C1 and C2) offer novel biological insights but are the most demanding in terms of analysis. They require dynamic PET data with sufficient temporal resolution, arterial or image-derived input function, and two-tissue compartment modelling (Volpi et al., 2023). The computational pipeline is more complex than other approaches, including voxel-wise parameter estimation, Laplace transform solutions and Euclidean distance-based similarity calculations. Despite these challenges, kinetic approaches provide unique information about metabolic dynamics that cannot be captured by static or tracer-based distribution methods.

fMRI offers the advantage of being non-invasive, widely available, and requiring no radiation exposure. Practically, fMRI acquisition is standard across many research settings, and the analytic pipelines is well-established and relatively computationally efficient. Because of these considerations, it was included here as a useful comparison to the PET-derived metrics. However, fMRI’s BOLD signal is complex and multi-faceted, reflecting a mixture of neural, vascular, and metabolic processes, and is subject to vascular confounds (e.g., neurovascular coupling), which can be especially prevalent in ageing (Deery et al., 2025b; Jamadar et al., 2025; Sala et al., 2023). While it measures a fundamentally related but different underlying neurobiology to FDG-PET, fMRI’s strong structural alignment make it valuable for multi-modal integration studies.

The results of this study should be interpreted in the context of its design and limitations. Firstly, the data was acquired under 50/50 bolus+infusion radiotracer administration, which enables the application of the fPET method. All other connectivity methods are usually applied to bolus-only PET acquisitions. Bolus+infusion protocols aim to maintain a constant plasma supply of radiotracer throughout the scan, which is not present during bolus acquisitions (Jamadar et al., 2022; Rischka et al., 2018). While data for the non-fPET Connectivity approaches was handled as closely as possible as outlined in their original sources, it remains possible that the presence of the constant radiotracer supply shaped the results. Secondly, the sample size, whilst large for PET/MR and fPET studies, is modest by comparison to other connectivity-behaviour studies. Lastly, while we have estimated measurement validity by analysing the predictive utility of each method for age and cognition, it remains possible that different methods may better predict other phenotypic variability not tested here, such as neurodegenerative disease, psychiatric illness, sex, and other factors.

In conclusion, our findings underscore that multi-method approaches are needed for a comprehensive understanding of metabolic brain networks, and that the physiological specificity of a connectivity measure is paramount for interpreting its relationship to behaviour and cognition. The optimal approach depends on the research question, available resources, and practical constraints. fPET Connectivity is recommended for cognition and ageing studies where dynamic PET is feasible; Tracer Distribution for age classification and single-scan designs with careful interpretation; fMRI for multi-modal integration and structure-function studies; and Kinetic C1 and C2 approaches for novel biological insights into metabolic dynamics and brain physiology. Researchers should weigh the trade-offs between physiological specificity, analytical complexity, acquisition requirements, and interpretability when selecting a connectivity approach for their studies.

## Statements & Declarations

## Declaration of the use of AI

AI (or other tools) were used to check code, spelling and grammar and improve parts of the text.

## Data and Code Availability

All data needed to evaluate and reproduce the results in the paper are present in the paper and/or the Supplementary Materials. Code for the calculation of the Kinetic C1 and C2 connectivity matrices cannot be made available at this time due to funder restrictions. The data and other codes are available upon request from the corresponding author.

## Author Contributions

All authors contributed to the study conception and design. Material preparation, data collection and analysis were performed by Hamish Deery, Emma Liang, Shenyue Zhao and Rui Zheng. The first draft of the manuscript was written by Hamish Deery and Sharna Jamadar and all authors commented on previous versions of the manuscript. All authors read and approved the final manuscript.

## Ethics approval

Approval was granted by the Human Research Ethics Committee of Monash University.

## Consent to participate

Written informed consent was obtained from the parents.

## Acknowledgements

We thank Robert Di Paolo, Gerard Murray, M. Navyaan Siddiqui, Katharina Voigt, Richard McIntyre, Lauren Hudswell and the staff at Monash Biomedical Imaging for their contributions to data acquisition and image reconstruction.

Code used in the calculation of Kinetic C1 and C2 connectivity was developed by and provided to us by Alessandra Bertoldo, from the University of Padua. We gratefully acknowledge this contribution.

The authors acknowledge the facilities and scientific and technical assistance of the National Imaging Facility, a National Collaborative Research Infrastructure Strategy (NCRIS) capability, at Monash Biomedical Imaging, Monash University.

## Supplementary Information

## 1. Participants demographics and cognition

The characteristics of the whole sample (N =80), as well as the younger (N = 36) and older (N = 44) participants, are shown in Table S1. The mean age of the whole sample was 54.5 years (SD = 24.5). The proportion of women was 53%. The average years of education was 17. Average BMI was 24.8 kg/m^2^, resting heart rate was 78.1 BPM and systolic and diastolic blood pressure were 136.2 and 78.1 mmHg, respectively. The mean fasting blood glucose was 5.0 mmol/L.

The mean age of the younger group was 28.4 years and the older group 75.8 years. The proportion of women in the younger group (52%) and older group (48%) was not significantly different. The average years of education was higher in the younger (18.1) than the older group (16.4).

The older group had significantly higher mean systolic blood pressure than the younger group (149.2mmHg vs 120.3mmHg). The older group also had a higher fasting blood glucose level (5.2 vs 4.8 mmol/L). The older group also has significantly worse performance on the HVLT (7.4 vs 9.2), category switch reaction time (2.11 vs 1.42s), stop signal RT (0.59 vs 0.53s), and digit symbol substitution (28.1 vs 64.5) performance.

**Table S1.** Demographics for the whole sample and comparison of older and younger groups. Continuous variables are mean (standard deviation); categorical variables are percentage.

|  | Whole sample (N = 80) |  | Younger (N = 36) |  | Older (N = 44) |  | p-FDR |
| --- | --- | --- | --- | --- | --- | --- | --- |
|  | Mean | SD | Mean | SD | Mean | SD |  |
| Age (years) | 54.5 | 24.5 | 28.4 | 6.1 | 75.8 | 6.1 | 0.000 |
| Sex (% Female) | 50% |  | 52% |  | 48% |  | 0.659 |
| Education (years) | 17.1 | 3.5 | 18.1 | 2.8 | 16.4 | 3.9 | 0.051 |
| Fasting blood glucose (mmol/L) | 5.0 | 0.5 | 4.8 | 0.4 | 5.2 | 0.6 | 0.004 |
| Systolic blood pressure (mmHg) | 136.2 | 25.7 | 120.3 | 17.3 | 149.2 | 24.3 | 0.000 |
| Diastolic blood pressure (mmHg) | 81.5 | 12.5 | 78.8 | 13.0 | 83.7 | 11.8 | 0.093 |
| Resting heart rate (bpm) | 78.1 | 15.6 | 82.7 | 17.3 | 74.4 | 13.2 | 0.029 |
| Body Mass Index (kg/m<sup>2</sup>) | 24.8 | 3.7 | 23.8 | 3.7 | 25.6 | 3.5 | 0.040 |
| HVLT: Delayed recall | 8.2 | 2.6 | 9.2 | 2.5 | 7.4 | 2.4 | 0.003 |
| Digit Symbol Substitution: Correct count | 12.1 | 2.2 | 12.3 | 2.7 | 12.0 | 1.8 | 0.595 |
| Category switch: RT in Switch Trials (sec) | 1.80 | 0.64 | 1.42 | 0.40 | 2.11 | 0.63 | 0.000 |
| Digit symbol substitution (Correc) | 44.2 | 22.5 | 64.5 | 12.2 | 28.1 | 14.1 | 0.000 |
| Stop signal: RT in stop signal trials | 0.56 | 0.13 | 0.53 | 0.13 | 0.59 | 0.12 | 0.029 |

<sup>†</sup>P-values are based on *t*-test for continuous (2-sided) and Ch-square for categorical variables.

## 2. Cognitive tests

### Hopkins Verbal Learning Test (HVLT)

The HVLT is a three-trial list learning and free recall task. The learning trials comprised 12 words, four words from each of three semantic categories. Approximately 20–25 minutes after the learning trials, participants completed delayed recall and recognition trials. The delayed recall required free recall of any of the 12 words. The recognition trial comprised 24 words, including the 12 target words and 12 false-positives, six semantically related, and six semantically unrelated. Delayed recall was calculated as the total words recalled.

### Task Switching

For task switching, a computerised test was used in which participants were presented with a word and had to perform a categorisation task. The categorisation task was dependant on two cues that appeared on screen across the trials. One cue was a heart symbol, for which participants were asked to categorise the word presented via a key press as either a LIVING or a NON-LIVING object. If the cue was an arrow-cross, participants were asked to categorise the word as either BIGGER or SMALLER than a basketball. The cue was randomised across trials. Half the trials were switch trials and half were non-switch trials. Half the switch and non-switch trials was congruent in the key presses for either task, half was incongruent. The task switching measure was latency of correctly responding to a switch trial.

### Stop Signal

The stop signal trial was a computer-based test in which participants were required to press the left response key if an arrow on screen pointed left and the right response key if the arrow pointed right. If a signal beep sounded, participants were instructed to stop their response. The delay between presentation of an arrow and signal beep started at 250ms and was altered up or down by 50ms based on performance. The delay increased up to 1150ms if the previous stop signal trial was successful and decreased to 50ms if the previous trial was unsuccessful. The stimulus onset asynchrony between the onset of a fixation circle at the start of each trial was 2000ms. Reaction time in the stop signal trials was recorded.

### Digit Symbol Substitution

A computer-based task presenting participants with a matrix of 18 column and 16 rows. Participants were required to translate symbols shown above the matrix in a key into digits in the matrix. The trial lasted two minutes. Performance was measured as total count of correct responses.

### Digit Span

A measure of verbal short term and working memory used in two formats: Forward and backward digit span. Participants were presented with a series of digits and are asked to repeat them in either the order presented (forward span) or in reverse order (backwards span). After two consecutive failures of the same length, the test was stopped. Scores were derived as the length of longest correct series for forward and backward recall.

## 3. Neuroimaging acquisition parameters and pre-processing

### Acquisition parameters

Participants were positioned supine in the scanner bore with their head in a 32-channel radiofrequency head coil. Anatomical T1- and T2-weighted MRI scans were acquired during the first 12 minutes. The T1 3D MPRAGE sequence parameters were: TR = 1,640 ms, TE = 2.34 ms, flip angle = 8°, field of view = 256 × 256 mm², voxel size = 1.0 mm isotropic, 176 slices, sagittal acquisition. For the T2 FLAIR sequence: TR = 5,000 ms, TE = 396 ms, field of view = 250 × 250 mm², voxel size = 0.5 × 0.5 × 1.0 mm³, 160 slices. List-mode PET and T2* EPI BOLD-fMRI sequences began 13 minutes into the scan. PET parameters were: voxel size = 1.39 × 1.39 × 5.0 mm³. EPI BOLD-fMRI parameters were: TR = 1,000 ms, TE = 39 ms, FOV = 210 mm², voxel size = 2.4 mm isotropic, 64 slices, ascending axial acquisition, total acquisition time = 40 minutes. A 40-minute resting-state scan was undertaken while participants viewed a movie of a drone flying over the Hawaiian Islands. At 58 minutes, diffusion-weighted imaging (DWI) was acquired to index white matter connectivity using a spin-echo sequence with the following parameters: 58 axial slices, 2.5-mm slice thickness, right-to-left phase encoding, 64 diffusion directions at b = 3,000 s/mm2, TR = 6,800 ms, TE = 171 ms, and a simultaneous multi-scale factor of 2. A single non-diffusion-weighted volume (b = 0 s/mm2) was additionally acquired with left-to-right phase encoding.

### Pre-processing

Participants’ PET data were binned into 344 3D sinogram frames (16 seconds per frame), corrected for attenuation (Burgos et al., 2014), and reconstructed using the Ordinary Poisson-Ordered Subset Expectation Maximization algorithm with point spread function correction (3 iterations, 21 subsets). Reconstructed DICOM slices were converted to NIFTI format (344 × 344 × 127 voxels; voxel size: 1.39 × 1.39 × 2.03 mm³) and a single 4D NIFTI volume constructed from the 3D volumes. The 4D PET volumes were motion-corrected (Jenkinson et al., 2002) and corrected for partial volume effects using a 25% gray matter threshold and surface-based spatial smoothing with a Gaussian kernel (FWHM = 8 mm) (Greve et al., 2016; Greve et al., 2014). Similarly, the T2* images underwent brain extracted, unwarping and motion correction using six rotation and translation parameters. The images were also temporally detrended, smoothed at 8mm FWHM, and normalised to MNI space. Finally, the FDG and fMRI time-series for each subject were denoised using anatomical component-based noise correction (aCompCor) (Whitfield-Gabrieli & Nieto-Castanon, 2012), For fPET, this produces a residual metabolic connectivity timeseries that measure the temporal dynamics of the glucose signal and removes the FDG uptake trend (Deery, Liang, Siddiqui, et al., 2024; Muschelli et al., 2014; Reed, Ponce de Leon, et al., 2025). The fMRI timeseries was also bandpass filtered at 0.01 Hz to 0.1 Hz (Whitfield-Gabrieli & Nieto-Castanon, 2012).

Structural Connectivity (SC) was assessed from diffusion-weighted imaging (DWI) data (Tournier et al., 2019). Fiber orientation distributions were estimated using single-shell multi-tissue constrained spherical deconvolution (CSD) (Dhollander, 2016). Whole-brain probabilistic tractography was then performed to reconstruct the white matter pathways, together with anatomically constrained tractography (ACT) (Smith et al., 2012; Yeh, 2018). For each participant, 22 million streamlines were generated, with streamline weights estimated using spherically informed filtering of tractograms (SIFT2) algorithm (Smith et al., 2015; Yeh, 2018).

## 4. Distance-based connectivity methods

### Kinetic C1 and C2

To obtain physiologically meaningful kinetic connectivity measures, we extracted the TACs from the dynamic FDG PET data, following the approach developed by Castellaro and colleagues (Catsellaro et al., 2017) and described by Volpi et al. (2023) for application to connectivity analyses. Dynamic PET data were fitted using Sokoloff’s two-tissue, three-rate-constant model, parameterised by four values: K₁ (tracer inflow, mL/cm³/min), k₂ (efflux, min⁻¹), k₃ (phosphorylation, min⁻¹), and Vb (blood volume fraction). The fitted rate constants define the net irreversible uptake macro-parameter K_i_ = K₁k₃ / (k₂ + k₃). Voxel-wise parameter estimation was performed using Variational Bayesian inference with cluster-based priors, as described elsewhere (Castellaro et al., 2017), using an image-derived input function (IDIF) as a surrogate for the plasma concentration Cp(t).

Following Volpi et al. (2023), the fitted total tissue curve Ct(t) was decomposed into two additive time courses. Adding the standard Sokoloff solutions for the first and second compartments and rearranging gives:

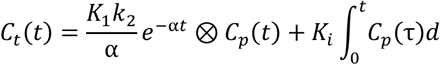

where α = k₂ + k₃ and ⊗ denotes convolution. The first term is an exponential-convolution component that decays with the effective washout rate α; the second is a cumulative-integral component that grows with the net uptake macro-parameter K_i_. These two curves sum to the tissue curve and provide complementary time courses that separate delivery-related and trapping-related information, echoing the C₁ vs C₂ distinction of Sokoloff’s model.

### Terminology note

We emphasise that these decomposed curves are not the exact compartment concentrations C₁(t) (free intracellular FDG) and C₂(t) (trapped FDG-6-phosphate) of the Sokoloff model. They are a convenient rewrite of the total tissue curve that captures the delivery-like and trapping-like behaviour of the tracer without explicitly solving for the two compartments. We therefore refer to them throughout as the delivery-like and trapping-like components of Ct, and we interpret the trapping-like component as the closest available proxy for phosphorylation-related signal, without claiming it is literally hexokinase activity.

Within-individual metabolic connectivity (wi-MC) was computed from region-level PET time courses using the Euclidean similarity metric, following Volpi et al. (2023). For each subject, three time series were extracted per region: (i) the full tissue time-activity curve (TAC) from the dynamic PET data; (ii) the delivery-like component of the fitted tissue curve; and (iii) the trapping-like component of the fitted tissue curve. The latter two are the additive decomposition of the fitted total tissue curve Ct described above.

For each pair of regions i and j, Euclidean similarity was computed as:

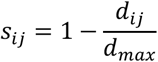

where d_ij_ is the Euclidean distance between the two time courses and d_max_ is the maximum Euclidean distance across all region pairs in that subject. Similarity values were then Fisher-z-transformed and rescaled to (0, 1) as described in Volpi et al. (2023). Matrices were computed separately for the full TAC, the delivery-like component, and the trapping-like component, then averaged across subjects to yield three group-level wi-MC matrices.

The two component-based matrices capture distinct physiological regimes: the delivery-like matrix emphasises tracer exchange at the blood–brain barrier (analogous to the early, flow-weighted portion of the tissue curve), while the trapping-like matrix emphasises net irreversible uptake and the trapping-related signal (analogous to the late, metabolism-weighted portion).

### Tracer Distribution Connectivity

For each subject, dynamic PET volumes were averaged to create a static 3D FDG uptake image. The Schaefer100 atlas was then applied to extract all voxel intensities within each of the 100 regions, preserving the full distribution of uptake values rather than collapsing to a single mean (Labarthe et al., 2026). For each subject and region, this yielded a variable-length vector of voxel intensities representing the regional tracer uptake distribution.

To reduce computational complexity while preserving the heavy tails of the SUV distributions (where critical pathological signals may reside), we applied a spatially-aware iterative compression algorithm from Labarthe et al. (Labarthe et al., 2026). Unlike uniform random down sampling, this algorithm adaptively merges voxels based on both spatial proximity and intensity similarity, ensuring that the resulting reduced feature set faithfully preserves the original metabolic topology and distributional shape.

Connectivity between regions was quantified using a distance-based approach combining energy distance and mutual information correction. For each pair of regions i and j, the distance between their voxel intensity distributions P and Q was computed using the energy distance metric:

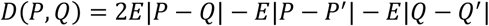

where P and Q are the voxel intensity distributions of two regions, and P’ and Q’ are independent copies. This metric measures the dissimilarity between distributions and is zero if and only if the distributions are identical. The energy distance was selected as it generalises the usual mean-based approach while remaining robust to noise in the tails of the distributions.

To account for non-linear statistical dependencies between regional uptake distributions, mutual information (MI) was incorporated into the distance calculation:

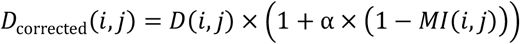

where α = 0.5 is a weighting parameter controlling the influence of mutual information on the distance correction. The MI quantifies the statistical dependence between two distributions, capturing both linear and nonlinear relationships. When SUV distributions are independent, MI is null and the distance is not affected; when a high dependency exists, the distance is scaled down, emphasising strong non-linear dependencies. The final connectivity strength between regions was defined using exponential similarity:

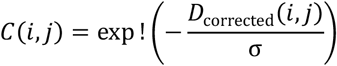

where σ = 1.0 is a scaling parameter. This exponential mapping emphasizes local neighbourhood structures through its rapid decay for increasing distances, effectively filtering out irrelevant distant connections. The resulting connectivity matrices capture the similarity of tracer uptake distributions across brain regions at the individual level, with the mutual information correction ensuring that regions with non-linear relationships are appropriately weighted.

## 5. Cost-efficiency analysis

We performed a cost-efficiency analysis at the 16-network level (8 networks × 2 hemispheres) and 100 region levels, testing sparsity levels from 5% to 90% in 5% increments to identify the optimal network density for each connectivity approach (Figure S1). Global efficiency was computed on binarised networks as the average inverse shortest path length between all pairs of nodes and cost as the proportion of connections retained at each threshold (Bullmore & Sporns, 2009). Cost-efficiency was then calculated as the difference between global efficiency and cost, with the optimal sparsity identified as the threshold maximising this value for each approach.

At the network level, optimal sparsity for Kinetic C2 and Tracer Distribution peaked at 10% sparsity, while fPET, fMRI, and Metabolic Covariance optimised at 15% sparsity, with an overall average of 13%. Metabolic connectivity and fMRI showed comparable cost-efficiency (∼0.64-0.69) and optimal sparsity (15%), suggesting these approaches capture similar network efficiency characteristics. At the region level, fPET, fMRI and Tracer Distribution peaked at 25% sparsity. Together, these results suggest that the 10-15% sparsity range consistently maximised cost-efficiency across all approaches at the network level, supporting the use of 15% sparsity threshold for subsequent analyses in the main manuscript. We also provide additional analyses at the 25% and 50% sparsity in Supplement 5.

**Fig S1.**
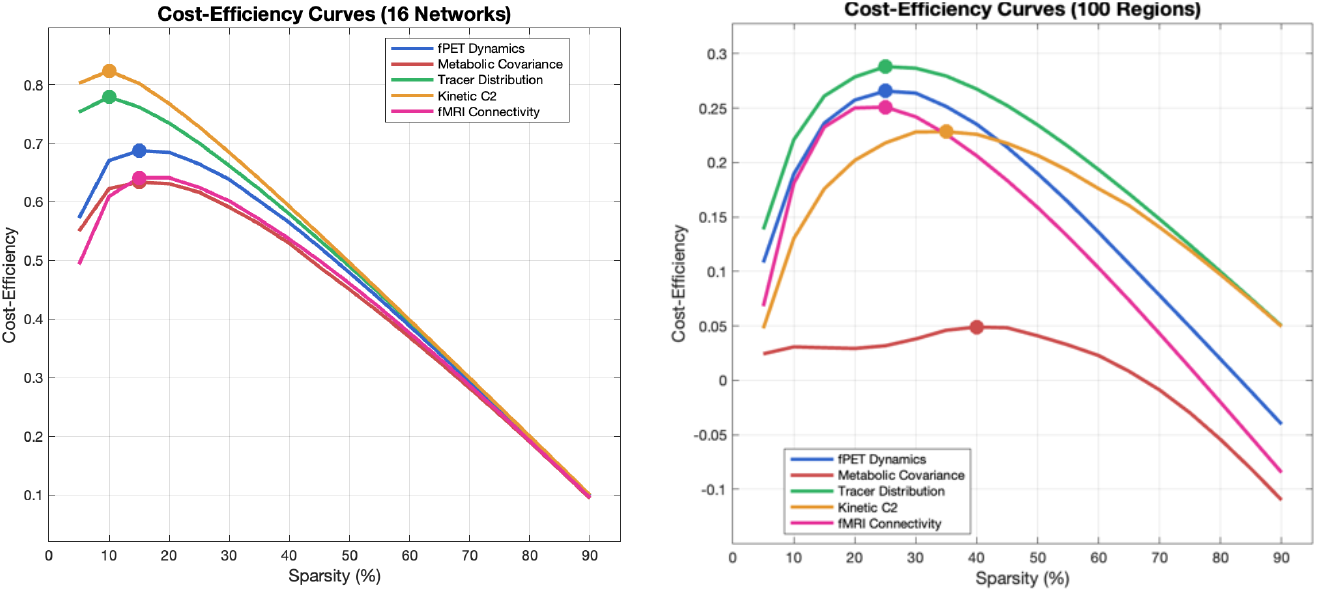
Cost-efficiency analysis. Sparsity analysis results at levels from 5% to 90% in 5% increments to identify the optimal density for the analysis of the connectivity approaches.

## 6. Graph metric definition

Network topology was characterised using four graph metrics computed on binarised networks (Bullmore & Sporns, 2009; Sporns & Betzel, 2016)::

### Clustering coefficient (C)

The average fraction of a node’s neighbours that are also connected to each other, reflecting local network segregation. For binary networks, this was computed as:

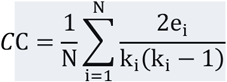

where *e_i_* is the number of edges between the neighbours of node i, *k_i_* is its degree, and *N* is the number of nodes.

### Characteristic path length (L)

The average shortest path length between all pairs of nodes, reflecting global network integration:

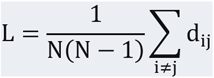

where *d_ij_* is the shortest path length between nodes i and j.

### Small-worldness (σ)

The ratio of normalised clustering to normalised path length where *C*_rand_ and *L*_rand_ are averages over 20 degree-matched random networks (10 rewiring iterations each). Values greater than 1 indicate small-world organisation.

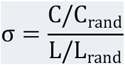

### Rich-club coefficient (Φ)

The degree to which high-degree nodes (hubs) are more densely interconnected than expected by chance. Computed as:

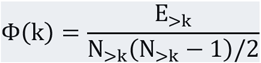

where *E*_>k_ is the number of edges among nodes with degree greater than k, and *N*_>k_ is the number of such nodes.

## 7. Graph metrics by connectivity approach

**Table S2.** Graph metric values (mean and SD) across connectivity approaches.

|  | Small worldness |  | Clustering coefficient |  | Path length |  | Rich club coefficient |  |
| --- | --- | --- | --- | --- | --- | --- | --- | --- |
|  | Mean | SD | Mean | SD | Mean | SD | Mean | SD |
| Metabolic Cov. (LOO) | 1.14 | 0.26 | 0.56 | 0.06 | 1.96 | 0.73 | 0.57 | 0.16 |
| fPET Connectivity | 1.00 | 0.03 | 0.41 | 0.07 | 2.17 | 0.23 | 0.43 | 0.12 |
| Tracer Distribution | 0.96 | 0.02 | 0.61 | 0.02 | 2.88 | 0.18 | 0.28 | 0.03 |
| Kinetic C1 | 0.73 | 0.07 | 0.71 | 0.05 | 4.25 | 0.61 | 0.25 | 0.05 |
| Kinetic C2 | 1.00 | 0.00 | 0.54 | 0.03 | 1.69 | 0.03 | 0.60 | 0.05 |
| fMRI Connectivity | 0.99 | 0.03 | 0.57 | 0.03 | 2.59 | 0.20 | 0.24 | 0.06 |

## 8. Gene expression results

**Table S4.**
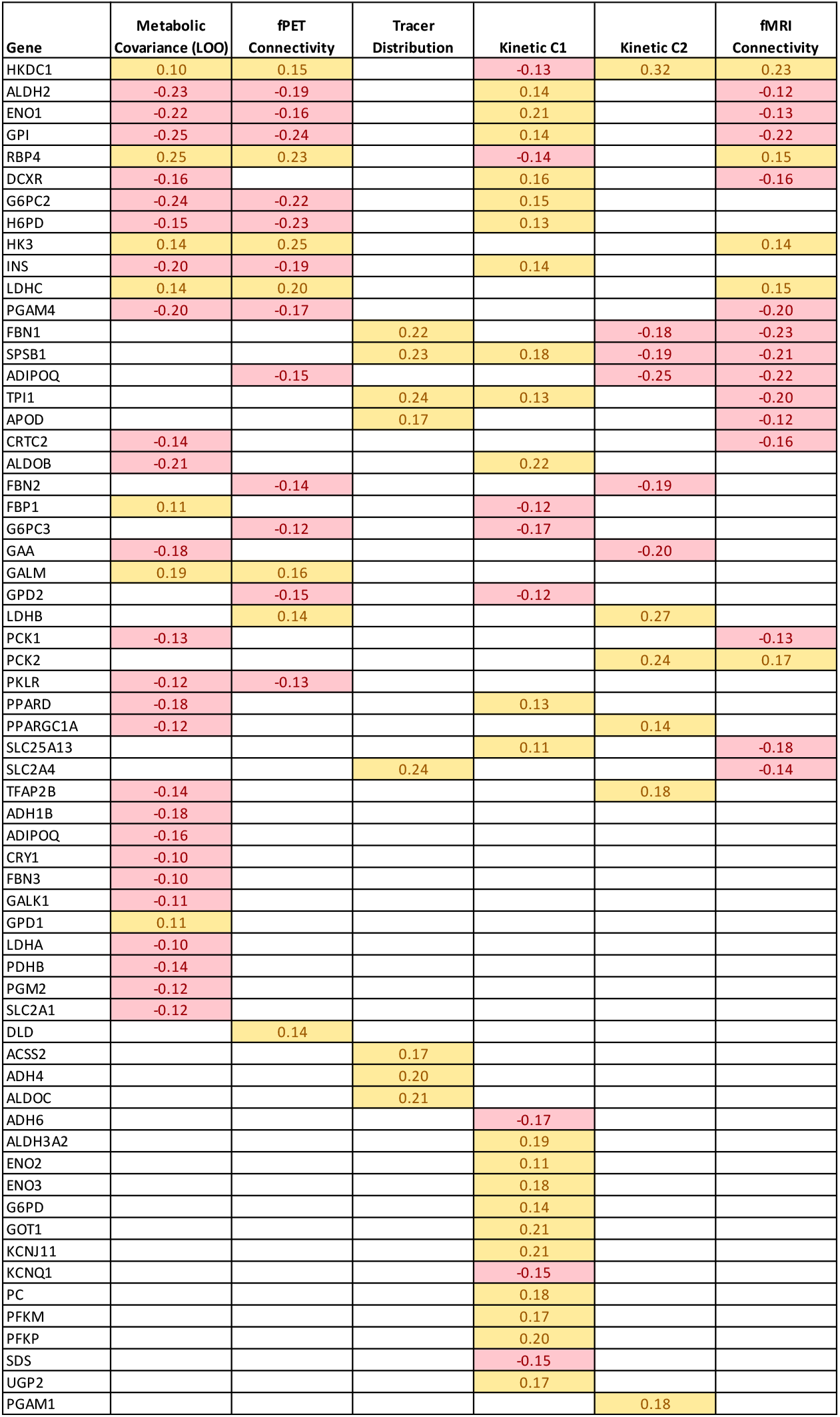
Significant genes (bootstrap 95% CI not crossing zero) for each connectivity approach. Genes are sorted by the number of approaches in which they were significant, from most-shared (top) to approach-unique (bottom). Yellow indicates positive PLSC gene loadings (higher gene expression associated with higher degree centrality); red indicates negative loadings. Blank cells indicate genes that were not reliably loaded in that approach. Numbers in cells are the PLSC loading values.

**Table S5.** Gene enrichment analysis. The significant genes from each approach (see Table S4) were subjected to pathway enrichment analysis in Metascape, with the understanding that the input gene list is pre-selected for glucose metabolism and enrichment results are therefore for descriptive purposes. *Count* is the number of input genes belonging to the given ontology term. *%* is the percentage of the approach’s significant genes that belong to that term. *Log10(q)* is the multi-test adjusted (FDR) p-value expressed in log base 10; more negative values indicate stronger enrichment (e.g., −10 corresponds to q = 10⁻¹⁰). Terms shown are those with a minimum count of 3 genes, an enrichment factor > 1.5, and an FDR-adjusted q < 0.05.

| Description | Count | % | Log10(q) |
| --- | --- | --- | --- |
| **Metabolic Covariance (LOO)** |  |  |  |
| Carbohydrate metabolic process | 27 | 84.38 | -43.28 |
| Glycolysis / gluconeogenesis | 17 | 53.12 | -33.82 |
| Metabolism of carbohydrates and carbohydrate derivatives | 15 | 46.88 | -18.15 |
| Gluconeogenesis | 10 | 31.25 | -18.03 |
| Glucagon signaling pathway | 10 | 31.25 | -13.93 |
| Small molecule catabolic process | 13 | 40.62 | -13.92 |
| Glucose homeostasis | 11 | 34.38 | -13.4 |
| Regulation of small molecule metabolic process | 9 | 28.12 | -7.96 |
| AMPK signaling pathway | 7 | 21.88 | -7.92 |
| Aerobic glycolysis augmented | 4 | 12.5 | -6.77 |
| Regulation of hormone levels | 8 | 25 | -5.15 |
| Regulation of nucleotide metabolic process | 4 | 12.5 | -3.47 |
| Diseases of carbohydrate metabolism | 3 | 9.38 | -3.03 |
| Alcohol metabolic process | 5 | 15.62 | -2.96 |
| Response to nutrient levels | 6 | 18.75 | -2.94 |
| Amino acid metabolism | 3 | 9.38 | -1.84 |
| Regulation of immune effector process | 4 | 12.5 | -1.32 |
| **fPET Connectivity** |  |  |  |
| Description | Count | % | Log10(q) |
| Glycolysis / gluconeogenesis | 13 | 68.42 | -26.37 |
| Glucose metabolic process | 12 | 63.16 | -22.32 |
| Pyruvate metabolic process | 9 | 47.37 | -16.51 |
| Glucose homeostasis | 7 | 36.84 | -8.14 |
| HIF-1 signaling pathway | 6 | 31.58 | -7.71 |
| Type II diabetes mellitus | 5 | 26.32 | -7.53 |
| Pyruvate metabolism | 5 | 26.32 | -7.41 |
| Glucagon signaling pathway | 5 | 26.32 | -5.84 |
| Alcohol metabolic process | 3 | 15.79 | -1 |
| Diseases of metabolism | 3 | 15.79 | -0.96 |
| Positive regulation of cell motility | 3 | 15.79 | -0.04 |
| **Tracer Distribution** |  |  |  |
| Description | Count | % | Log10(q) |
| Glycolysis / gluconeogenesis | 4 | 50 | -4.52 |
| Glucose metabolic process | 4 | 50 | -4.4 |
| Glycolysis and gluconeogenesis | 3 | 37.5 | -3.31 |
| **Kinetic C1** |  |  |  |
| Description | Count | % | Log10(q) |
| Glucose metabolic process | 23 | 76.67 | -49.61 |
| Glycolysis / Gluconeogenesis | 15 | 50 | -29.04 |
| Glycerol metabolism including linked IMDs | 8 | 26.67 | -15.18 |
| Glucose 6-phosphate metabolic process | 6 | 20 | -10.68 |
| Galactose metabolism | 6 | 20 | -9.89 |
| Alcohol metabolic process | 9 | 30 | -9.17 |
| Cori cycle | 5 | 16.67 | -8.98 |
| Regulation of hormone levels | 8 | 26.67 | -5.34 |
| NAD+ metabolic process | 4 | 13.33 | -4.77 |
| Pyruvate metabolism | 4 | 13.33 | -4.42 |
| Regulation of small molecule metabolic process | 6 | 20 | -3.82 |
| Response to xenobiotic stimulus | 6 | 20 | -3.67 |
| NADP+ metabolic process | 3 | 10 | -3.09 |
| Diseases of carbohydrate metabolism | 3 | 10 | -3.02 |
| Response to organophosphorus | 4 | 13.33 | -3.01 |
| miR targeted genes in muscle cell | 4 | 13.33 | -1.3 |
| **Kinetic C2** |  |  |  |
| Description | Count | % | Log10(q) |
| Glucose metabolic process | 9 | 81.82 | -17.14 |
| Glucose homeostasis | 6 | 54.55 | -6.99 |
| Pyruvate metabolic process | 4 | 36.36 | -5.04 |
| Glucagon signaling pathway | 4 | 36.36 | -4.06 |
| Fat cell differentiation | 3 | 27.27 | -2.55 |
| Eye morphogenesis | 3 | 27.27 | -2.32 |
| **fMRI Connectivity** |  |  |  |
| Description | Count | % | Log10(q) |
| Glucose metabolic process | 14 | 73.68 | -27.91 |
| Glycolysis / Gluconeogenesis | 10 | 52.63 | -18.33 |
| Gluconeogenesis | 8 | 42.11 | -14.89 |
| Glucose homeostasis | 9 | 47.37 | -11.75 |
| Small molecule catabolic process | 8 | 42.11 | -8.07 |
| Glucagon signaling pathway | 5 | 26.32 | -5.79 |
| Type II diabetes mellitus | 4 | 21.05 | -5.26 |
| Cori cycle | 3 | 15.79 | -4.46 |
| Response to xenobiotic stimulus | 4 | 21.05 | -1.98 |
| Regulation of lipid metabolic process | 3 | 15.79 | -0.75 |

## 9. Sensitivity analysis at 25% sparsity

To assess the robustness of our findings across topological scales, we repeated the analyses from the main manuscript at a higher sparsity threshold of 25% (Supplementary Tables S4–S10). The core conclusions from the 15% main analyses were largely preserved. First, the differential topological organization of connectivity approaches remained stable: Metabolic Covariance (both LOO and group-average) and fPET Connectivity continued to exhibit the strongest small-world properties, while Kinetic C1 consistently lacked small-world characteristics due to its prolonged characteristic path lengths.

Second, the cognitive associations showed remarkable consistency – fPET Connectivity remained the only approach with significant simple correlations (clustering: r = 0.48, rich-club: r = 0.34), and ridge regression confirmed that age dominates cognitive variance across all approaches, with R² values increasing from ∼0.02–0.21 without age to ∼0.42–0.44 when age was included.

Third, the structural connectivity comparisons were highly concordant: fMRI Connectivity and Metabolic Connectivity showed the strongest edge overlap with structural connectivity (DICE = 0.52 and 0.47, respectively), while Kinetic C2 remained the outlier with the poorest overlap (DICE = 0.18) and the most negative edge-strength correlations (ρ = -0.68). Finally, the genetic associations were similarly preserved, with Metabolic Covariance approaches continued to show the strongest PLSC correlations (r = 0.71, 36 significant genes), followed by Tracer Distribution (r = 0.581), fMRI (r = 0.534), and fPET (r = 0.512), while Kinetic C1 and C2 remained non-significant.

Despite this overall consistency, several differences in the gene-connectivity associations at 25% sparsity warrant mention. Metabolic Covariance (LOO) and fPET Connectivity showed robust positive associations at both thresholds, and their gene loading vectors remained strongly convergent (r = 0.77 at 25% vs. r = 0.81 at 15%). Kinetic C1 and Kinetic C2 showed robust negative associations at both thresholds, consistent with their anti-correlation with the metabolic approaches at 15% (Kinetic C1 vs. Metabolic Covariance (LOO): r = −0.33 at 15%, r = −0.45 at 25%). Tracer Distribution and fMRI Connectivity showed sign reversal in their degree–gene associations between the two thresholds.

The strongest cross-approach convergence between Metabolic Covariance (LOO) and fPET Connectivity (r = 0.81 at 15%, r = 0.77 at 25%) was robust across sparsity levels. The anti-correlation between Metabolic Covariance (LOO) and Kinetic C1 also strengthened at 25% (from r = -0.33 to r = -0.45), reinforcing the biological dissociation between the metabolic and kinetic approaches.

The results for Tracer Distribution and fMRI Connectivity suggests these approaches may capture a composite of multiple biological processes whose relative weighting shifts as the connectivity threshold changes. This contrasts with the Metabolic Covariance (LOO), fPET Connectivity and Kinetic C1 and C2 approaches, whose associations were stable across sparsity levels and likely reflect a single dominant biological signal. Tracer Distribution’s 25% signature in particular showed a mix of gluconeogenic and glycolytic loadings, distinguishing it from the metabolic approaches’ more unified glycolytic signature. This may reflect the unique biological basis of voxel-level FDG distribution patterns relative to other connectivity estimation methods.

Taken together, these findings confirm that our main conclusions are robust across sparsity levels that metabolic-based approaches (particularly fPET Connectivity) are most tightly coupled to cognition, structural connectivity, and genetic architecture, while kinetic approaches show minimal or opposite associations. The 25% results provide complementary insights into how different connectivity methods capture age-related and topological reorganisation at coarser network scales, and reveal which approaches have largely sparsity-stable biology versus threshold-dependent biology.

**Table S6.** Descriptive statistics for graph metrics of connectivity matrices at 25% sparsity.

|  | Small worldness |  | Clustering |  | Path length |  | Rich club |  |
| --- | --- | --- | --- | --- | --- | --- | --- | --- |
|  | Mean | SD | Mean | SD | Mean | SD | Mean | SD |
| fPET Connectivity | 1.00 | 0.01 | 0.50 | 0.07 | 1.92 | 0.15 | 0.41 | 0.11 |
| Metabolic Cov (LOO) | 0.99 | 0.20 | 0.68 | 0.05 | 2.25 | 0.61 | 0.53 | 0.17 |
| Tracer Distribution | 0.98 | 0.01 | 0.68 | 0.02 | 2.29 | 0.09 | 0.34 | 0.03 |
| Kinetic C1 | 0.85 | 0.05 | 0.75 | 0.03 | 2.95 | 0.31 | 0.33 | 0.06 |
| Kinetic C2 | 1.00 | 0.00 | 0.64 | 0.02 | 1.60 | 0.03 | 0.60 | 0.04 |
| fMRI Connectivity | 1.00 | 0.01 | 0.61 | 0.03 | 2.01 | 0.09 | 0.28 | 0.03 |
| Metabolic Cov. (Group Avg) | 0.98 | NA | 0.65 | NA | 2.29 | NA | 0.42 | NA |

**Table S7.**
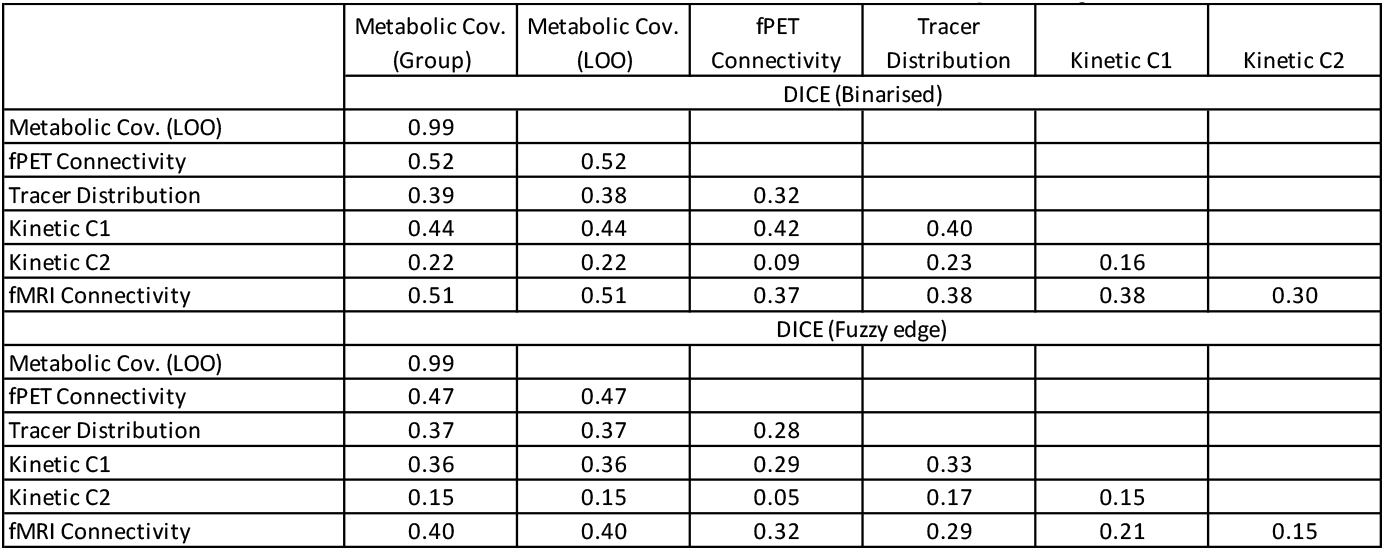
Dice coefficients of connectomes at 25% sparsity.

**Table S8.** Age group differences in network topology at 25% sparsity.

|  | Small worldness |  |  | Clustering |  |  | Path length |  |  | Rich club |  |  |
| --- | --- | --- | --- | --- | --- | --- | --- | --- | --- | --- | --- | --- |
|  | t-value | p-FDR | Cohens' d | t-value | p-FDR | Cohens' d | t-value | p-FDR | Cohens' d | t-value | p-FDR | Cohens' d |
| fPET Connectivity | -0.18 | 0.96 | -0.04 | 5.56 | 0.00 | 1.25 | 2.09 | 0.12 | 0.47 | 3.08 | 0.01 | 0.69 |
| Metabolic Cov. (LOO) | -0.08 | 0.96 | -0.02 | -2.19 | 0.05 | -0.49 | -0.72 | 0.71 | -0.16 | 0.84 | 0.40 | 0.19 |
| Tracer Distribution | -1.41 | 0.49 | -0.32 | 2.20 | 0.05 | 0.49 | -1.64 | 0.21 | -0.37 | -7.88 | 0.00 | -1.77 |
| Kinetic C1 | -0.22 | 0.96 | -0.05 | 0.80 | 0.43 | 0.18 | -0.51 | 0.74 | -0.11 | -1.76 | 0.12 | -0.39 |
| Kinetic C2 | 0.04 | 0.96 | 0.01 | 0.90 | 0.43 | 0.20 | 0.07 | 0.95 | 0.01 | -2.10 | 0.08 | -0.47 |
| fMRI Connectivity | 1.81 | 0.45 | 0.41 | -3.36 | 0.00 | -0.75 | -2.84 | 0.03 | -0.64 | -1.64 | 0.13 | -0.37 |

**Table S9.** Cognition PC associations with network topology at 25% sparsity.

|  | Small worldness |  | Clustering |  | Path length |  | Rich club |  |
| --- | --- | --- | --- | --- | --- | --- | --- | --- |
|  | Correlation | p-FDR | Correlation | p-FDR | Correlation | p-FDR | Correlation | p-FDR |
| fPET Connectivity | 0.00 | 0.999 | 0.48 | 0.00 | 0.21 | 0.25 | 0.34 | 0.01 |
| Metabolic Cov. (LOO) | 0.02 | 0.999 | -0.26 | 0.06 | -0.08 | 0.57 | 0.03 | 0.90 |
| Tracer Distribution | 0.07 | 0.999 | -0.05 | 0.79 | -0.17 | 0.29 | -0.40 | 0.00 |
| Kinetic C1 | 0.11 | 0.999 | -0.09 | 0.64 | -0.14 | 0.32 | 0.07 | 0.84 |
| Kinetic C2 | -0.03 | 0.999 | 0.03 | 0.81 | 0.04 | 0.71 | 0.01 | 0.90 |
| fMRI Connectivity | 0.14 | 0.999 | -0.22 | 0.09 | -0.19 | 0.25 | -0.23 | 0.07 |

**Table S10.** Summary of structural connectivity analyses at 25% sparsity.

|  | DICE (binary) | Edge Pearson r | EdgeSpearman p | Network DICE | Network Corr | Hub Ratio | Hub overlap | Prediction R<sup>2</sup> | p-value |
| --- | --- | --- | --- | --- | --- | --- | --- | --- | --- |
| fPET Connectivity | 0.47 | 0.16 | -0.13 | 0.91 | 0.34 | 1.40 | 16.00 | 0.03 | 0.00 |
| Metabolic Cov. (LOO) | 0.48 | 0.04 | -0.10 | 0.85 | 0.17 | 1.28 | 12.00 | 0.00 | 0.15 |
| Tracer Distribution | 0.37 | -0.17 | -0.33 | 0.87 | 0.20 | 0.86 | 8.00 | 0.03 | 0.00 |
| Kinetic C1 | 0.32 | -0.14 | -0.40 | 0.84 | -0.08 | 1.22 | 8.00 | 0.02 | 0.00 |
| Kinetic C2 | 0.18 | -0.39 | -0.68 | 0.79 | -0.20 | 0.26 | 0.00 | 0.14 | 0.00 |
| fMRI Connectivity | 0.52 | 0.27 | 0.04 | 0.86 | 0.40 | 1.16 | 4.00 | 0.07 | 0.00 |

**Table S11.**
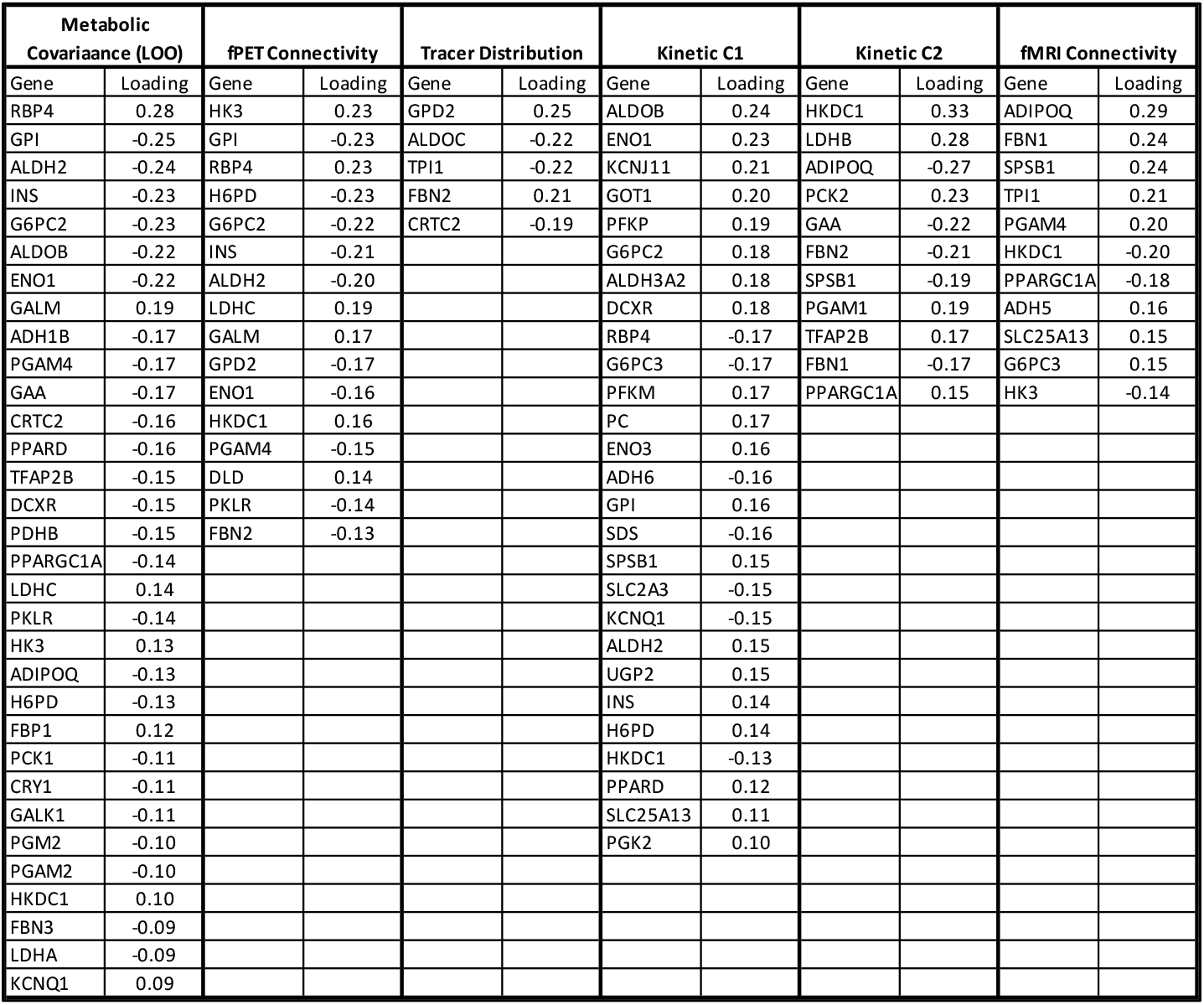
Significant genes (bootstrap 95% CI not crossing zero) for the PLSC associations between gene expression and network degree at 25% sparsity.

